# The skin commensal yeast *Malassezia* promotes tissue homeostasis via the aryl hydrocarbon receptor

**DOI:** 10.1101/2025.03.26.645515

**Authors:** Eduardo Gushiken-Ibañez, Michelle Stokmaier, Giuseppe Barone, Alessia Staropoli, Tugay Karakaya, Dietmar Beer, Francesco Vinale, Giuseppe Ianiri, Salomé LeibundGut-Landmann

## Abstract

As an abundant fungal colonizer of human skin, *Malassezia* has long been associated with pathological skin conditions, yet its role in skin homeostasis remain poorly understood. Here, we demonstrate that *Malassezia furfur* plays an active role in maintaining epidermal integrity by producing tryptophan-derived metabolites that activate the aryl hydrocarbon receptor (AhR), a key regulator of keratinocyte differentiation and inflammation. Using a fungal mutant defective in indole production, we show that *M. furfur*-derived AhR activation is required to restore barrier function and control inflammation in diseased skin. AhR-deficient mice fail to benefit from *M. furfur*-mediated barrier protection, underscoring the importance of microbial-derived AhR agonists in skin physiology. These findings establish a previously unrecognized mutualistic role for *Malassezia* in epidermal homeostasis, challenging its perception as solely a pathogenic fungus and expanding our understanding of the skin microbiota’s influence on barrier function and immune regulation.

**KEY FINDINGS:**

- *Malassezia*-derived indoles reprogram epidermal gene expression to enhance keratinocyte function.
- AhR activation by *Malassezia* restores skin barrier integrity and reduces inflammation.
- *Malassezia* Sul1-dependent tryptophan metabolism is essential for the production of AhR agonists.
- The barrier protective effects of *Malassezia* are mediated specifically through keratinocyte intrinsic AhR signaling.

## INTRODUCTION

The skin functions as a vital barrier separating the body from the external environment and protecting it against physical, chemical, and microbial threats [1, 2]. Central to this function is the structural and functional organization of the epidermis, a stratified epithelium made of keratinocytes. The stratum corneum, the outermost layer of the skin epidermis consisting of terminally differentiated keratinocytes, known as corneocytes and embedded within a lipid-rich matrix, limits water permeability and shields the body from harmful substances [3]. This physical and biological barrier is defective in various common skin disorders, including atopic dermatitis, psoriasis, and ichthyosis [3–5], which are in turn characterized by increased transepidermal water loss (TEWL), sensitivity to environmental irritants, and susceptibility to infections [6–8]. Therefore, preserving the integrity of the epidermis is critical for maintaining tissue integrity and homeostasis of the host. The processes governing skin barrier restoration and maintenance are however not fully understood. Furthermore, the relative contributions of endogenous factors vs. exogenous influences including microbial metabolites to these processes remain unclear. Understanding these interactions is key towards identifying novel strategies that aim at restoring skin barrier function in common cutaneous pathologies.

The aryl hydrocarbon receptor (AhR) is an important regulator of skin homeostasis [9]. As a cytoplasmic receptor it senses diverse environmental cues in response to which it elicits a ligand-dependent response resulting in detoxification of xenobiotic compounds, modulation of host defenses and/or the regulation of epithelial cell differentiation and function [10].

Upon activation, AhR translocates to the nucleus, where it forms a heterodimer with the aryl hydrocarbon receptor nuclear translocator (ARNT) and binds DNA response elements. Target genes include the Cytochrome P450 1A1 gene (*CYP1A1*) encoding for the subunit A of the Cytochrome P450 multiprotein complex, which in turn metabolize xenobiotics and other potentially toxic compounds [11]. Importantly, AhR also regulates the expression of genes encoding proteins with structural roles in the skin such as Ovo like protein 1 (*OVOL1*), Filaggrin (*FGL*), Involucrin (*IVL*) or Loricrin (*LOR*) [12]. While the last 3 have direct roles in defining the structure of the skin, *OVOL1* is a transcription factor promoting the skin barrier function [13]. In keratinocytes, AhR signaling regulates the expression of structural skin barrier genes and genes involved keratinocyte differentiation [14–18]. Additionally, AhR activation modulates skin responses to oxidative stress, further underscoring its importance in maintaining epidermal integrity [19].

Members of the microbiota have been proposed to act as a source of indoles serving as AhR ligands, including *Staphylococcus spp.* [20], *Corynebacterium spp.* [21] and the yeast *Malassezia* [22–24]. By constituting >90% of the skin mycobiome, *Malassezia* spp. are the most abundant fungal colonizer of the human skin, thriving in the lipid-rich environments due to their unique dependence on host-derived lipids for survival [25]. *M. restricta* and *M. globosa* are most abundant on human skin, followed by *M. sympodialis* and *M. furfur* [26]. Unlike many other fungi, *Malassezia* lacks the ability to synthesize fatty acids de novo, making it highly adapted to sebaceous regions of the skin such as the scalp, face, and upper trunk, where sebum production is highest [27, 28]. Despite its commensal lifestyle, *Malassezia* has been associated with certain skin disorders, such as atopic dermatitis, seborrheic dermatitis, and pityriasis versicolor (PV) [29], all of which are characterized by a disrupted skin barrier and chronic inflammation. While topical antifungal treatments are sometimes used for atopic dermatitis, their effectiveness remains inconclusive [30]. This lack of consistent efficacy raises the possibility that *Malassezia* is not the primary causative agent but rather an opportunistic factor that exacerbates symptoms in individuals with a pre-existing impaired skin barrier. Thus, the exact role of *Malassezia* in these conditions remains unclear. Instead of directly triggering disease, *Malassezia* may take advantage of a weakened barrier and contribute to worsening inflammation, suggesting that barrier dysfunction itself plays a crucial role in disease development.

*Malassezia* is known to produce bioactive brown pigments by metabolizing tryptophan, which is naturally present in the skin as a component of sweat [31, 32]. *M. furfur*, in particular, generates indolic compounds that can bind and activate AhR [22, 24, 33]. The biological relevance of these tryptophan-derived fungal metabolites remains however unknown. The prevalence of *Malassezia* on human skin raises the question whether *Malassezia*-derived indoles play a role in shaping skin physiological processes. Understanding these interactions is also important given the association of *Malassezia* and common pathological skin conditions [34], which leaves room for speculation whether fungal metabolites may favor the development of disease, or rather prevent disease and actively support skin health.

Here, we show that *Malassezia*-derived indoles activate AhR signaling in keratinocytes, leading to the modulation of keratinocyte function. We further show that activation of AhR by the fungal metabolites restored the barrier function of disrupted and inflamed skin. This effect was mediated by keratinocyte intrinsic AhR activation and lost with *Malassezia* mutant strains unable to produce AhR-activating indoles. The ability of *Malassezia* to produce AhR agonists emphasizes its role as a mediator of skin homeostasis and provides an important example of how the abundant commensal yeast provides a benefit to the host.

## RESULTS

### *M. furfur*-derived metabolites induce strong AhR activation via production of indoles in a tryptophan dependent manner

To assess the ability of *Malassezia* to metabolize tryptophan, we evaluated two different strains of *M. furfur* under three distinct growth conditions including Tween 80 agar [33], a modified mDixon medium without mycological peptone (mDix-mp) [35], and a defined minimal medium (MM) supplemented with increasing concentrations L-tryptophan. *M. furfur* strain CBS 1878, previously reported to produce pigments from tryptophan but characterized by aneuploidy [33, 35], was compared to CBS 141414, a haploid strain with a high-quality genome assembly [36] and greater amenability to genetic manipulation [37, 38], but with an unknown capacity for pigment production. After 4 and 8 days of incubation, both CBS 1878 and CBS 141414 produced brown pigments both in MM agar and mDix-mp agar (**Figure 1A-B**), although on MM agar, CBS 1878 exhibited better growth, and a higher degree of pigment production compared to CBS 141414 at both time points (**Figure 1B**). On Tween 80 agar, only *M. furfur* CBS 1878 displayed robust growth and strong production of brown pigments (**Supplementary Figure S1A**). Regardless of the growth medium and strain, pigment intensity correlated with increasing tryptophan concentrations, indicating a dose-dependent relationship between tryptophan availability and pigment biosynthesis. The pigments were observed as both intracellularly absorbed, resulting in brown-colored cells, and extracellularly secreted into the medium, suggesting an active process of pigment production and release.

**Figure 1.**
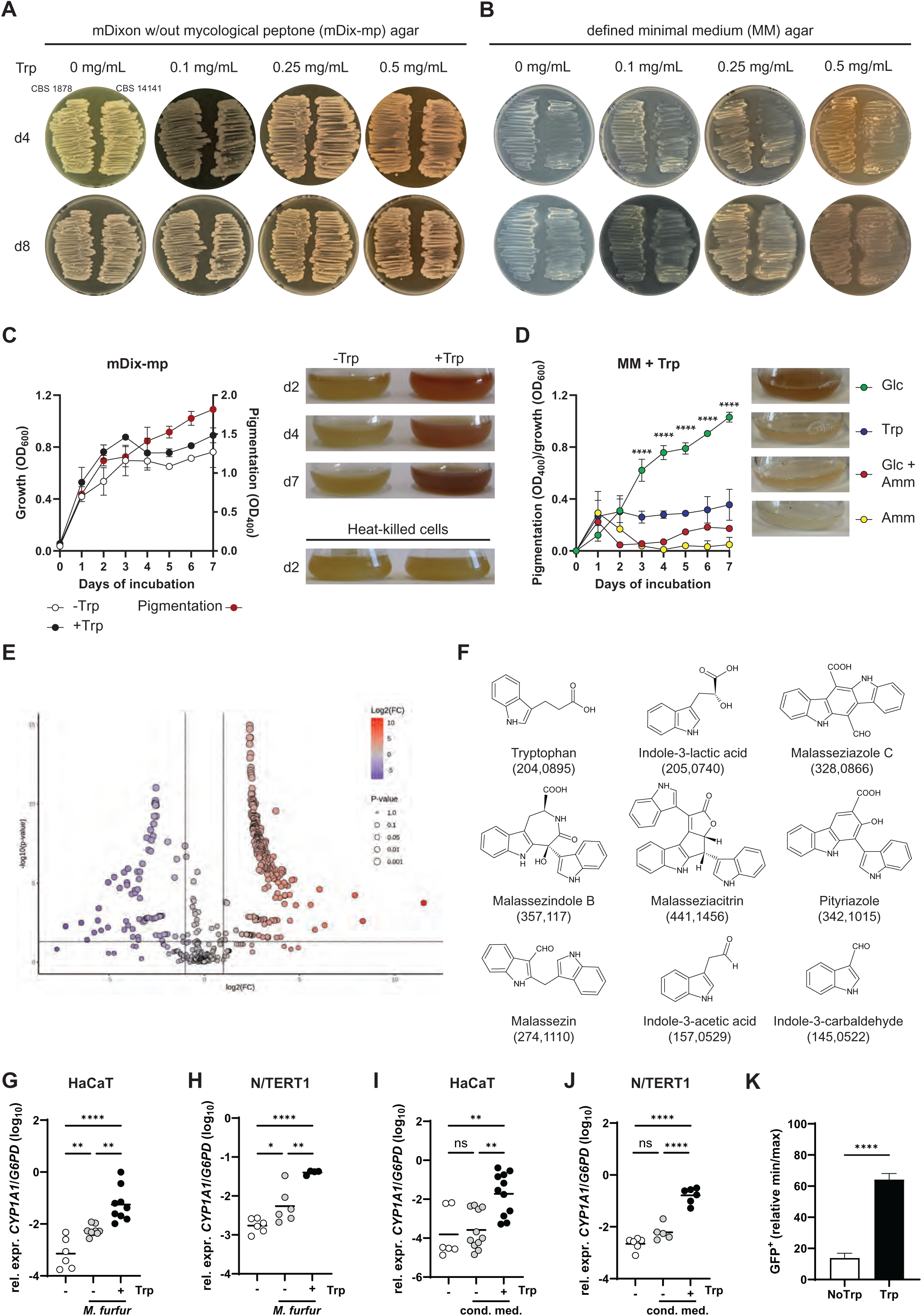
**M. furfur-derived metabolites induces strong AhR activation via production of indoles in a tryptophan dependent manner.** A. -B. *M. furfur* strains CBS 1878 and CBS 14141 grown for 4 and 8 days on mDix-mp agar (A) or on MM agar (B) supplemented with 0, 0.1, 0.25 and 0.5 mg/mL of L-tryptophan. C. *M. furfur* CBS 14141 growth (OD_600_, left axis) and pigment release (OD_400_, right axis) over 7 days incubation in liquid mDix-mp in presence or absence of L-tryptophan. D. *M. furfur* CBS 14141 pigment production relative to the growth (OD_400_/OD_600_) in liquied MM supplemented with tryptophan and glucose as carbon source (Glc), ammonium nitrate as nitrogen source (Amm), glucose and ammonium nitrate (Glc + Amm), or with tryptophan as sole carbon and nitrogen source (Trp). E. – F. Untargeted metabolomics of *M. furfur* CBS 14141 grown in mDix-mp supplemented with or without tryptophan. Volcano plot of differentially produced metabolites by *M. furfur* when grown in presence of tryptophan (E) and structure of detected indoles produced by *M. furfur* (F). G. – J. *CYP1A1* expression by HaCaT (C, E) and N/TERT1 (D, F) keratinocyte cell lines after 24 hours of infection with *M. furfur* (C, D) or exposure to conditioned medium of *M. furfur (E, F)* grown in presence (+) or absence (-) of tryptophan. Unstimulated cells were included as a control (open symbols). n = 4 – 11 / group pooled from 2 - 5 independent experiments. The mean of each group is indicated. K. GFP expression by the reporter cell line H1G1-1C3 in response to 24 hours of stimulation with conditioned medium from *M. furfur* (I, J) grown in presence (+) or absence (-) of tryptophan. Expression levels are displayed relative to unstimulated and FICZ-stimulated cells. Bars are the mean + SEM of 9 separately stimulated wells pooled from 2 independent experiments. The statistical significance of differences between groups was determined by two-way ANOVA (D), one-way ANOVA (G-F) or unpaired *t*-test (G), * p < 0.05, ** p < 0.01, *** p < 0.001, **** p < 0.0001. See also Supplementary Figure S1.

Given the aim of generating *M. furfur* mutants with impaired ability to produce pigments, further experiments were carried out using only the haploid M*. furfur* strain CBS 14141 in liquid medium which allowed quantification of growth and pigment production by optical density at OD_600_ and OD_400_, respectively. In liquid mDix-mp supplemented with 0.5 mg/mL of tryptophan, the pigment production by *M. furfur* CBS 14141 was visible from the second day of incubation and increased over time, in contrast to cultures grown without tryptophan, which remained pale (**Figure 1C**). Heat-killed cells were unable to produce pigments, indicating that tryptophan degradation depended on the metabolic activity of the yeast (**Figure 1C**).

To determine whether *M. furfur* produces pigments when tryptophan serves as its sole nitrogen and/or carbon source, we supplemented MM broth in four different ways: (1) MM + glucose + tryptophan (tryptophan serving as the sole nitrogen source, glucose as the sole carbon source), (2) MM + ammonium nitrate + tryptophan (tryptophan as the carbon source, ammonium nitrate as the nitrogen source), (3) MM + tryptophan alone (tryptophan as both, the carbon and nitrogen source), and (4) MM + glucose + ammonium nitrate + tryptophan (both carbon and nitrogen sources present). Pigment production relative to growth, measured as OD_400_/OD_600_, was highest when *M. furfur* was grown in MM supplemented with glucose and tryptophan. Growth in MM with tryptophan alone or with glucose and ammonium nitrate resulted in similar but lower pigment levels. No pigments were detected when ammonium nitrate and tryptophan were provided together (**Figure 1D**). Similarly, *M. furfur* produced pigments on solid medium with Tween 80 serving as a carbon source and tryptophan a nitrogen source (**Figure S1A**). These findings indicate that pigment production requires not only tryptophan as a nitrogen source but also the presence of an additional carbon source, aligning with previous results from Mayser et al. [33].

Untargeted metabolomics was then carried out for *M. furfur* CBS 14141 grown in mDixon - mp in the presence or absence of tryptophan, revealing 339 significant molecular features (FDR < 0.05; log2FC ± 2), with 256 upregulated and 83 downregulated compounds (**Figure 1E**). Among the detected metabolites, eight were identified exclusively when *M. furfur* was grown in presence of tryptophan, including some that were previously described: indole-3-lactic acid, malasseziazole C, malassezindole B, indole-3-carbaldehyde, pityriazole, malassezin [39], indole-3-acetic acid, and malasseziacitrin [33, 40] (**Figure 1F**). A complete list of the detected metabolites is provided in **Supplementary Table S1**.

To examine whether the tryptophan metabolites released from *M. furfur* contained AhR agonists, we exposed HaCaT and N/TERT1 keratinocyte cell lines to *M. furfur* CBS 14141 grown in presence or absence of tryptophan and assessed activation of the AhR pathway based on induction of the proximal AhR target gene *CYP1A1* after 24 hours of stimulation. While *M. furfur* alone elicited a basal response independently of tryptophan, AhR activation was strongly enhanced when pigment production was induced by tryptophan (**Figure 1G-H**). To verify that the AhR-stimulating molecules were contained within the secretome of *M. furfur*, we incubated keratinocytes with conditioned medium prepared from *M. furfur* grown in presence of tryptophan and again observed robust AhR activation; conversely, the response to control conditioned medium from *M. furfur* grown in absence of tryptophan did not differ from that of untreated keratinocytes (**Figure 1I-J**). AhR activation by *M. furfur*-derived metabolites was also conserved in primary human keratinocytes (**Supplementary Figure S1B-C**). Our findings were further confirmed by quantifying AhR activation by means of an AhR reporter cell line expressing GFP under the control of AhR binding domain XRE (H1G1-1C3 [41]) (**Fig. 1K**). Together, these findings corroborate that *M. furfur*-derived soluble tryptophan metabolites are potent AhR agonists, directly binding to AhR and activating downstream signaling.

The capacity to activate AhR signaling through tryptophan-derived metabolites was not unique to *M. furfur*; however, *M. furfur* exhibited significantly stronger AhR activation compared to all other *Malassezia* species tested, including *M. pachydermatis*, *M. globosa*, *M. restricta*, and *M. sympodialis*, regardless of the tryptophan concentration used (**Supplementary Figure S1D, E**). While these other species induced only baseline levels of AhR activation, *M. furfur* uniquely stimulated a markedly higher response, as indicated by significantly increased GFP reporter activity. This distinction suggests that *M. furfur* possesses a enhanced metabolic capacity to generate potent AhR-activating metabolites compared to its relatives.

### Activation of AhR by *M. furfur* in the epidermis modulates the expression of structural protein encoding genes

We next aimed at understanding the functional consequences of *Malassezia*-induced AhR activation in the skin. As keratinocytes grown in medium-submerged monolayer cultures do not well reflect the situation in the epidermis made of a stratified epithelium, we generated organotypic human epidermal equivalents (HEEs) [42]. Consistent with what we observed in keratinocyte monolayer cultures, both tryptophan-processing fungal cells and soluble tryptophan metabolites strongly induced AhR signaling in HEEs as evidenced by pronounced upregulation of *CYP1A1* expression (**Supplementary Figure S2A-B**). Importantly, we also found genes encoding structural proteins critical for skin barrier function, upregulated (**Fig. 2A**). These included involucrin (*IVL*) and small proline-rich protein 2A (*SPRR2A*), which are essential components of the cornified envelope contributing to the mechanical resilience and permeability barrier of the skin [43–45], as well as the transcriptional regulator *OVOL1,* a key regulator of keratinocyte differentiation orchestrating the transition of basal into terminally differentiated keratinocytes essential for maintaining the stratified epidermal structure [46]. The induction of these genes aligned with the previously reported role of AhR in enhancing keratinocyte differentiation and enhancing barrier integrity [47]. The activity of *M. furfur* driving this response was contained within the fungal secretome, as confirmed by comparing stimulation of HEEs with soluble tryptophan metabolites to infection of HEEs with *M. furfur* (**Supplementary Figure S2C-D**).

**Figure 2.**
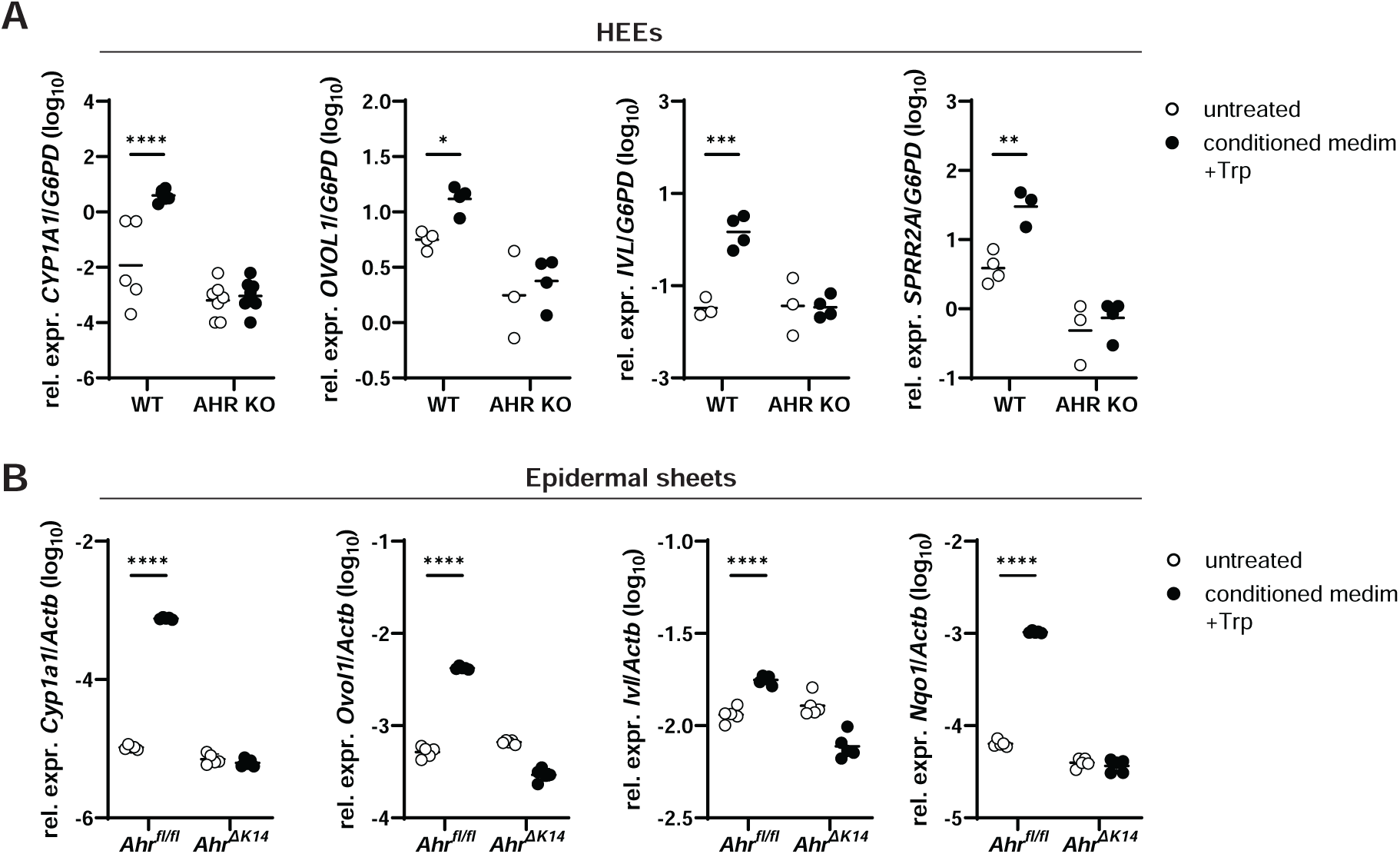
Activation of *AhR* by *M. furfur* in the epidermis modulates the expression of structural protein encoding genes. **A.** *CYP1A1*, *OVOL1*, *IVL*, and *SPRR2A* expression by HEEs made of WT or AhR-deficient N/TERT1 cells after 24 hours of stimulation with conditioned medium from *M. furfur* grown with tryptophan (filled symbols) or left untreated (open symbols). n = 5 - 7 / group pooled from 2 independent experiments (CYP1A1); n = 3 – 4 / group (other genes). **B**. *Cyp1a1*, *Ovol1*, *Ivl*, and *Nqo1* expression by epidermal sheets obtained from the ear skin of *Ahr^fl/fl^* or *Ahr*^Δ*K14*^ mice after 24 hours of incubation in conditioned medium from *M. furfur* grown with (filled black symbols) or without tryptophan (filled grey symbols) or left untreated (open symbols). n = 5 / group pooled from 2 independent experiments. The mean of each group is indicated. The statistical significance of differences between groups was determined by two-way ANOVA (A) or one-way ANOVA (B), * p < 0.05, ** p < 0.01, **** p < 0.0001. **See also Supplementary Figure S2.**

To confirm that the induction of skin barrier genes by the fungal metabolites was AhR-dependent in HEEs, we generated *AHR* knockout N/TERT1 cells by means of CRISPR/Cas9 gene editing (**Supplementary Figure S2A**) and generated HEEs from the knockout cells. As expected, *AHR*-deficient HEEs in comparison to the parental wildtype (WT) HEEs failed to induce *CYP1A1*, *OVOL1*, *IVL*, and *SPRR2A* expression in response to stimulation with *M. furfur*-derived tryptophan metabolites (**Figure 2A**).

To further reinforce these findings, we extended our experiments to epidermal sheets obtained from ear skin of naïve mice. Unlike the *in vitro* reconstructed HEE model, which relies on stratification under artificial culture conditions, epidermal sheets retain the original complex architecture of *bona fide* epidermis and contain immune cells, which are naturally hosted in the epidermis. Stimulation of epidermal sheets with *M. furfur* tryptophan metabolites stimulated the induction of *Ovol1* and *Ivl* (**Figure 2B**). We also found *Nqo1* upregulated in epidermal sheets stimulated with fungal metabolites (**Figure 2B**) reflecting activation of the Nrf2 pathway [48]. This indicated that the activation of the AhR pathway by *Malassezia* metabolites was conserved across different skin models. To confirm AhR-dependence and to confirm that keratinocytes were responsible for mediating AhR signaling in response to fungal metabolites in epidermal sheets, we repeated the experiment with epidermal sheets from *Ahr*^Δ*K14*^ mice lacking AhR selectively in keratinocytes. *Cyp1a1*, *Ovol1*, *Ivl*, and *Nqo1* expression was indeed abolished in the mutant epidermal sheets confirming that AhR signaling was keratinocyte-intrinsic (**Figure 2B**). Together, these results suggested that *M. furfur*-induced tryptophan metabolites have the capacity to modulate skin barrier architecture through AhR-dependent transcriptional regulation of structural protein encoding genes in keratinocytes.

### *M. furfur*-induced AhR signaling restores barrier integrity of inflamed skin

Based on our *in vitro* data, we wondered whether *M. furfur* would modulate skin barrier function *in vivo* by triggering AhR signaling in the epidermis. We further speculated that such an effect would be particularly relevant in the context of pathological skin conditions characterized by barrier defects, such as atopic dermatitis. We therefore applied mild tape stripping to disrupt the stratum corneum [49], thereby mimicking in mice one of the key hallmarks of atopic dermatitis [50] prior to associating the ear skin with *M. furfur* [51]. We first verified that AhR was activated in the murine skin in response to indole producing *M. furfur* (+Trp) (**Supplementary Figure S3A**) without affecting fungal colonization (**Supplementary Figure S3B**). As shown previously [52], colonization of barrier impaired murine skin with *M. furfur* led to exacerbated inflammation, as evaluated by measuring the increase in skin thickness, which reached a maximum between day 3 and day 5 after colonization (**Figure 3A**). In contrast, association of tape-stripped skin with *M. furfur* grown in presence of tryptophan to induce the secretion of AhR agonists suppressed the induction of inflammation as evidenced by reduced ear thickness (**Figure 3A**), reduced epidermal thickness (**Figure 3B-C**), and reduced neutrophil counts in the skin tissue (**Figure 3D-E**). Importantly, the anti-inflammatory effect of *M. furfur*-derived indoles was abolished in AhR-deficient mice (**Figure 3A-E**).

**Figure 3.**
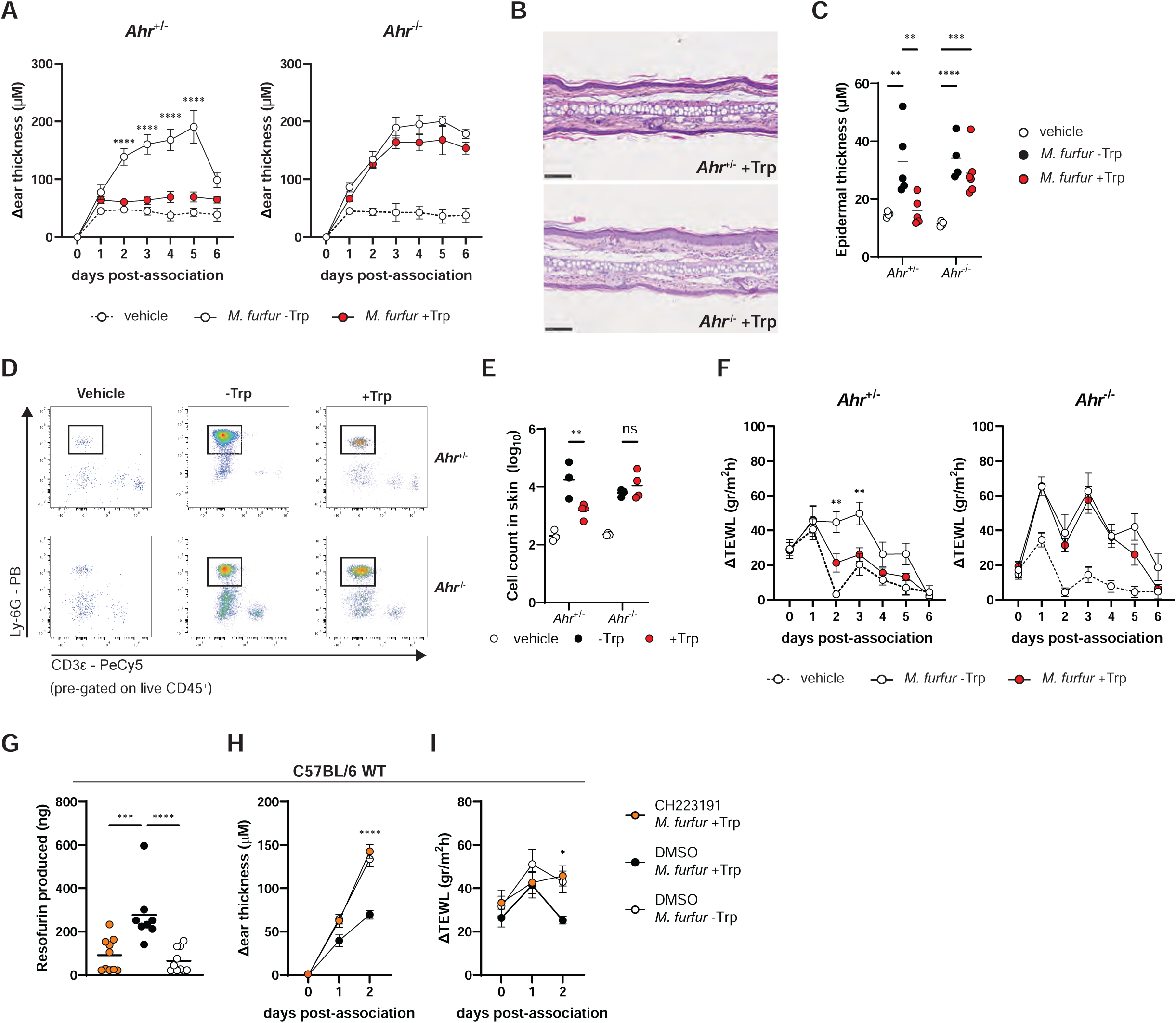
*M. furfur* induced AhR signaling restores barrier integrity in inflamed skin. The ear skin of *AhR^-/-^*mice and *AhR^+/-^* littermate control mice (A-F) or of WT C57BL/6 mice (G-I) was mildly tape stripped prior to epicutaneous administration of *M. furfur* grown in presence (+Trp) or absence of tryptophan (-Trp). A group of control mice was tape stripped and treated with olive oil (vehicle). **A.** Increase in skin thickness in *AhR^-/-^* (B) and *AhR^+/-^*littermate control mice (A) at the indicated time point relative to pre-treatment. n = 8 - 10 / group pooled from 2 independent experiments, except for the vehicle treated groups for which n = 4. The mean ± SEM is indicated. **B. – C.** Representative images of H&E-stained histology sections (B) and quantification of epidermal thickness (C) on day 7 after fungal association. n = 4 - 6 / group. Each symbol represents the average epidermal thickness per mouse. The mean of each group is indicated. **D. – E.** Neutrophils in the ear skin of colonized mice were quantified by flow cytometry 3 days after fungal association. Neutrophil counts per ear (D) and representative plots (E). n = 3 – 4 / group. Each symbol represents one mouse. **F.** TEWL in *AhR^-/-^* (F) and *AhR^+/-^* littermate control mice (E) at the indicated time points. n = 6 – 9 / group pooled from 2 independent experiments, except for the *AhR^-/-^* -Trp group where n = 4. The mean ± SEM is indicated. **G. – I.** Mice were treated with CH223191 or DMSO solvent prior to tape stripping and fungal association. Cyp1a1 activity on day 2 (G), increase in skin thickness relative to pre-treatment (H) and TEWL (I) at the indicated time points. n = 10 / group pooled from 2 independent experiments. Each symbol represents one mouse (G) or the mean ± SEM (H, I). The statistical significance of differences between the *M. furfur* +Trp and *M. furfur* -Trp groups for each mouse genotype (A, C, D, F) or between CH223191-treated vs. DMSO-treated groups associated with *M. furfur* +Trp (G-I) was determined by two-way ANOVA (A-F, H-I) or one-way ANOVA (G), * p < 0.05, ** p < 0.01, **** p < 0.0001. **See also Supplementary Figure S3.**

Barrier-impaired skin exhibits a defect in water retention, which results in increased trans-epidermal water loss (TEWL). As such, TEWL serves as a reliable and widely used functional parameter to evaluate skin barrier integrity [2]. We therefore assessed TEWL in the skin of *M.furfur*-associated animals and found that TEWL raised when barrier-impaired skin was colonized with *M. furfur* grown without tryptophan (**Figure 3F**), confirming that the fungus exacerbated skin barrier defect when not producing indoles. The degree of TEWL reached a maximum between two and four days after fungal association (**Figure 3F**). In contrast, colonization with indole producing *M. furfur* strongly impaired the increase in TEWL (**Figure 3F**), supporting a role of *Malassezia*-derived tryptophan metabolites in the restoration of the skin barrier. Of note, this skin protective effect of indole producing *M. furfur* was abolished in *Ahr*-deficient mice (**Figure 3F**). In *AhR^-/-^* mice, TEWL levels raised in response to tape stripping and *M. furfur* administration and then remained elevated, regardless of indole production profile of the fungus (**Figure 3F**), underscoring the critical role of AhR in the response to the tryptophan metabolites.

To exclude that the observed AhR-dependent phenotype resulted from pre-existing structural and/or functional perturbances in the epidermis due to environmental or dietary AhR ligands [14, 53], we used a pharmacological inhibitor of AhR signaling (CH223191) [54] in WT C57BL/6 mice. This approach allowed us to block AhR activity selectively during the duration of the experiment (**Figure 3G**) without interfering with AhR signaling during skin development. Consistent with our previous results obtained in genetically modified *AhR^-/-^* mice, treating WT mice with the inhibitor completely abolished the effect of fungal tryptophan metabolites. Skin thickness and TEWL levels in the skin of CH223191-treated mice colonized with indole producing *M. furfur* were as high as those in skin-associated with *M. furfur* not producing tryptophan metabolites (**Figure 3H-I**), confirming that the phenotype is not due to pre-existing structural differences in the skin or putative effects of diet.

Together, these results underscore the pivotal role of *Malassezia*-derived metabolites at the fungus-host interface. By preventing inflammation and restoring skin barrier integrity in an AhR-dependent manner, those *Malassezia*-derived metabolites mediate a host-protective effect during microbial colonization.

### AhR-mediated skin barrier restoration in response to *M. furfur* is keratinocyte intrinsic

AhR signaling is widely expressed across various cell subsets in the skin, including keratinocytes, fibroblasts, and immune cells, of which each has a different role in skin homeostasis. Our *in vitro* experiments with keratinocytes and HEEs, which revealed major transcriptional changes in skin barrier genes (**Figure 2**), suggested that keratinocytes may be primary targets of fungal AhR agonists in the skin. To confirm that the functional implications of AhR activation by *M. furfur*-derived tryptophan metabolites in the murine skin occurred keratinocyte-intrinsically, we repeated the skin colonization experiments in *Ahr*^Δ*K14*^ mice lacking AhR selectively in keratinocytes, while other cell types retained an intact *Ahr* gene (**Supplementary** Figure 4A). Fungal colonization of *Ahr*^Δ*K14*^ mice phenocopied largely what we observed in full-body AhR knockout mice. The association of barrier impaired ear skin with *M. furfur* grown in presence of tryptophan failed to improve skin barrier function (**Figure 4**) in contrast to what we observed in *AhR^fl/fl^*littermates, while fungal colonization was not affected (**Supplementary Figure S4B**). More specifically, the conditional knockout mice exhibited elevated inflammation (**Figure 4A**), epidermal thickness (**Figure 4B-C**) and TEWL (**Figure 4D**) in both tryptophan-treated and untreated cells compared to their cre-negative counterparts, in which barrier integrity was restored as evidenced by a limited skin thickness, inflammation and TEWL (**Figure 4A-D**). Together, these findings demonstrate that keratinocyte-intrinsic AhR signaling induced by indole producing *M. furfur* is essential for maintaining skin barrier integrity.

**Figure 4.**
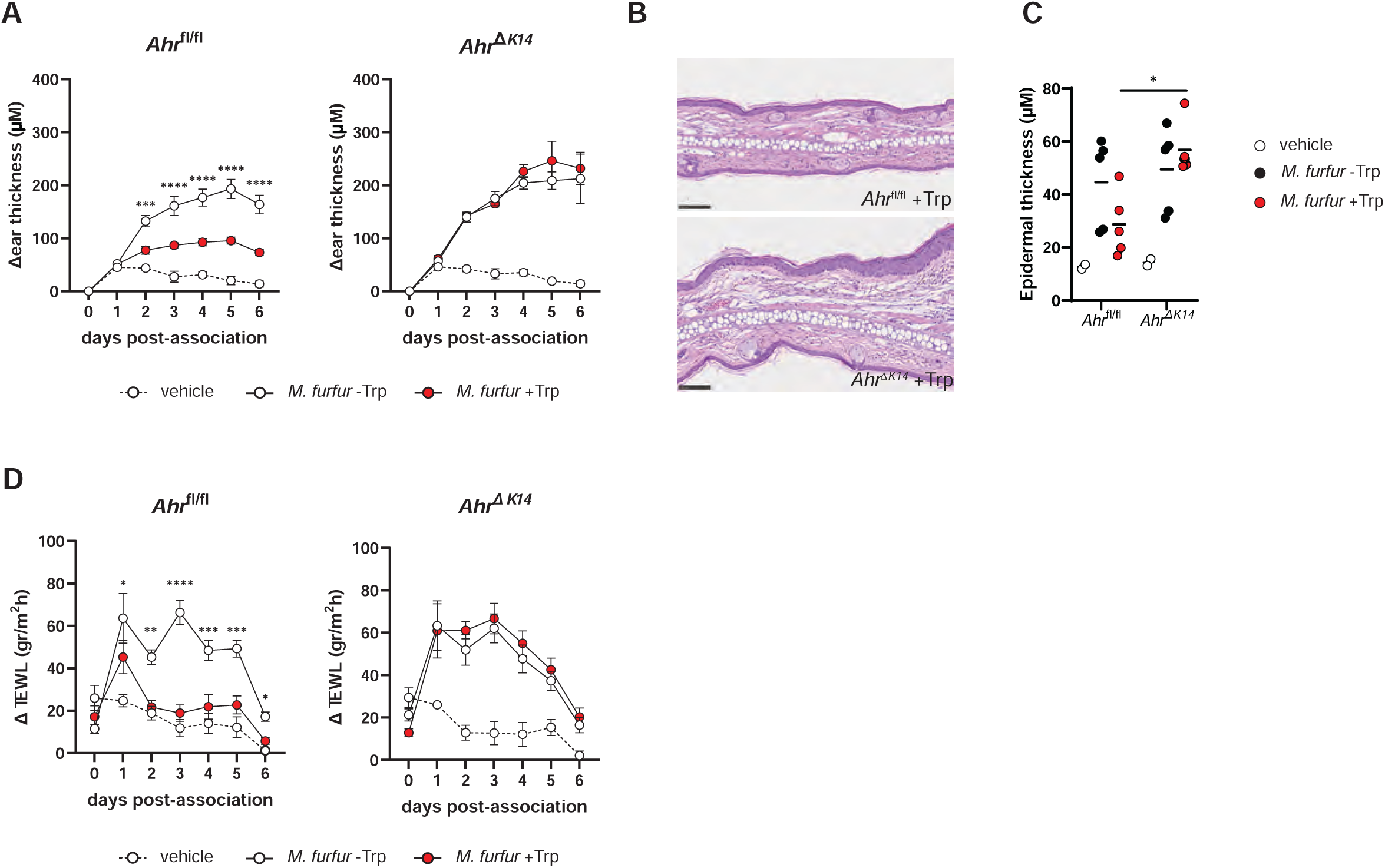
AhR-mediated skin barrier restoration in response to *M. furfur* is keratinocyte intrinsic. The ear skin of *Ahr*^Δ*K14*^ mice and *Ahr^fl/fl^* littermate control mice was mildly tape stripped prior to epicutaneous administration of *M. furfur* grown in presence (+Trp) or absence of tryptophan (-Trp). A group of control mice was tape stripped and treated with olive oil (vehicle). **A.** Increase in skin thickness in *AhR^-/-^* and *AhR^+/-^* littermate control mice at the indicated time point relative to pre-treatment. n = 8 / group pooled from 2 independent experiments, except for the vehicle group where n = 4 - 5. The mean ± SEM is indicated. **B. – C.** Representative images of H&E-stained histology sections (B) and quantification of epidermal thickness (C) on day 7 after fungal association. Each symbol represents the average epidermal thickness per mouse. The mean of each group is indicated **D.** TEWL in *AhR^-/-^* and *AhR^+/-^* littermate control mice at the indicated time points. n = 8 / group pooled from 2 independent experiments, except for the vehicle group where n = 4 - 5. The mean ± SEM is indicated. The statistical significance of differences between the *M. furfur* +Trp and *M.* furfur -Trp groups (A, D) or between genotypes (C) was determined by two-way ANOVA, * p < 0.05, ** p < 0.01, **** p < 0.0001. **See also Supplementary Figure S4.**

### A forward genetic screen identifies a *M. furfur sul1* mutant with impaired indole production

To identify *M. furfur* mutants defective in indole production, we generated an *Agrobacterium*-mediated random-insertional mutagenesis library and evaluated 600 individual transformants grown in mDix-mp in the presence of tryptophan assessing pigment loss in multiwell plates to prevent diffusion on agar plates. Visual inspection over the course of 12 days allowed the identification of a small number of *M. furfur* transformants displaying impaired pigment production (**Figure 5A**). Two transformatns (mutants 1G1 and 2B4) could be confirmed for their reduced ability to produce brown pigments compared to the parental WT when grown independently in a larger volume of medium (**Figure 5B**). While prolonged incubation of mutant 1G1 in presence of tryptophan resulted in the production of a red pigment, mutant 2B4 was profoundly impaired in pigment production and was therefore selected to be further investigated. Inverse PCR and amplicon sequencing confirmed a T-DNA insertion in a sulphate transporter (accession KAI3628368) (**Figure 5C**, top) that is closest ortholog of the *S. cerevisiae* gene *SUL1*, which is the gene name that we use in this work. Note that this gene is mistakenly named *SUL2* in Genbank under accession KAI36d8368. In the model yeast, Sul1 is a high affinity sulphate permease responsible for sulfate uptake and regulation of endogenous activated sulfate intermediates [55].

**Figure 5.**
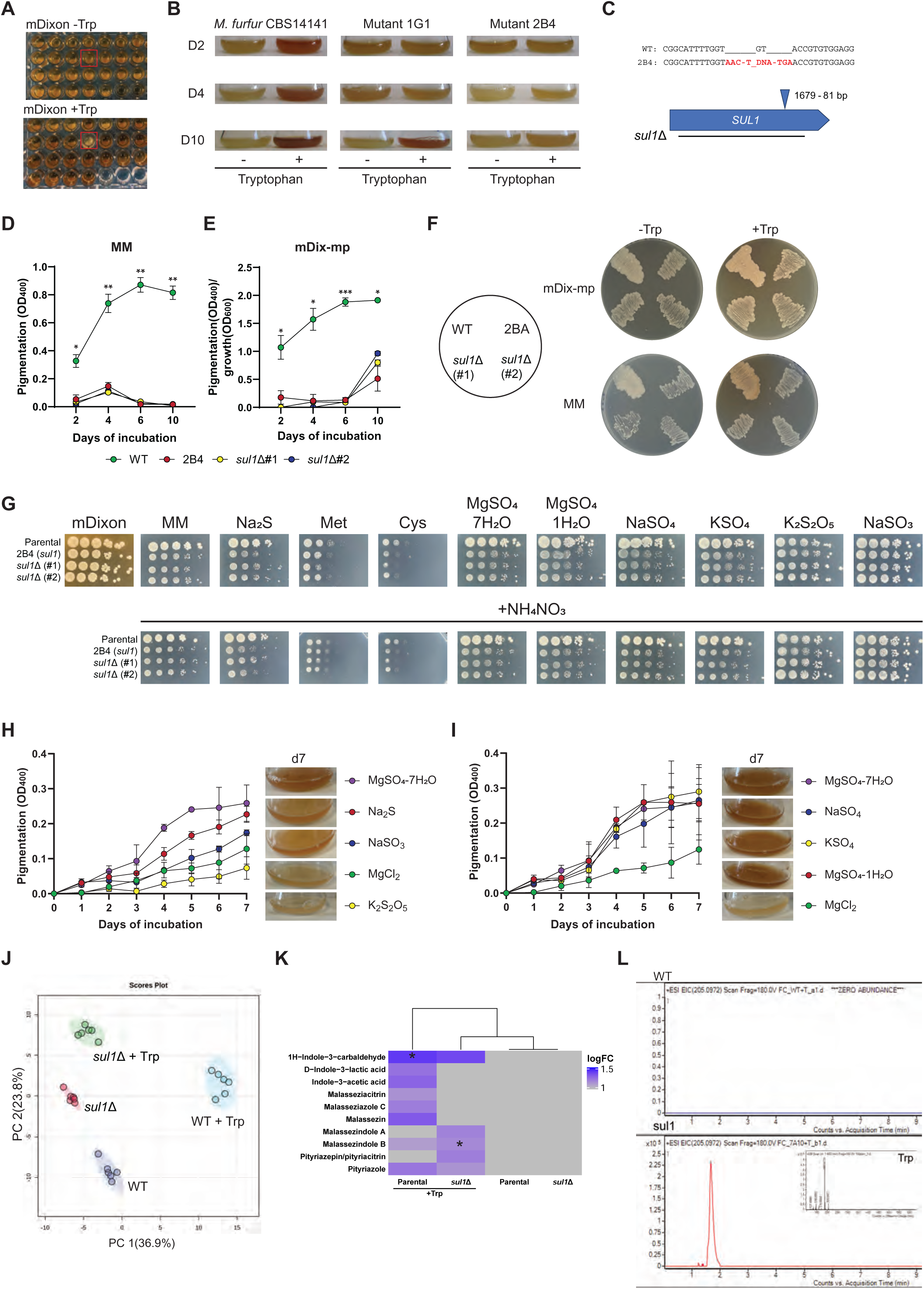
A forward genetic screen identifies a M. furfur sul1 mutant with impaired indole production. **A. – C.** Random insertional and targeted mutagenesis in *M. furfur* CBS 14141. Representative insert of a 96 well plate during the screening of random insertional mutants grown in mDix-mp supplemented without (bottom) or with (top) tryptophan (A). *M. furfur* CBS 14141 and two random insertional mutants 1G1 and 2B4 grown in mDix-mp supplemented with or without tryptophan for 2, 4 and 10 days (B). Schematic representation of *SUL1* mutagenesis (C). Top: T-DNA insertion site in the *SUL1* gene in strain 2B4 aligned with the WT sequence. T-DNA sequence in red, including the canonical TGA stop codon. **D. – F.** Pigmentation (OD_400_) and pigmentation relative to the growth (OD_400_/OD_600_) of *M. furfur* CBS14141, *sul1* random insertional mutant (2B4) and targeted mutant (*sul1*Δ#1 and *sul1*Δ#2) in liquid MM (D) and mDix-mp (E) supplemented with tryptophan. The same strains were also tested in mDix-mp agar and MM agar supplemented with or without tryptophan (F); the sheme indicates the position of the strains on the plates. **G – I.** Phenotype analysis of the *M. furfur* CBS 14141 and the strains 2B4 and *sul1*Δ#1 and *sul1*Δ#2 mutants on different inorganic and organic sulphate sources. MM agar in which MgSO_4_ was replaced with MgCl_2_ was supplemented with 2 mM of sodium sulfide (Na_2_S), potassium metabisulfite (K_2_S_2_O_5_), sodium sulfite (Na_2_SO_3_), magnesium sulphate (MgSO_4_ x 7 H_2_0, and MgSO_4_ x 1H_2_O), sodium sulphate (Na_2_SO_4_), and potassium sulphate (KSO_4_) (G). Pigmentation (OD_400_) of parental *M. furfur* CBS14141 in liquid MM supplemented with the same sulfur sources as in 5G (H-I). **J. – L.** Untargeted metabolomics of parental *M. furfur* (WT) and *sul1*Δ clone 1 supplemented with (+Trp) or without tryptophan. PCA plot of metabolomic data (J). Heatmap showing the abundance of *M. furfur*-produced indoles (K). EIC (extracted ion chromatogram) relative to tryptophan signal in parental *M. furfur* (WT) and *sul1*Δ1 mutant grown in presence of tryptophan (L). In D and E, the statistical significance of difference between the parental *M. furfur* (WT) and the *sul1* mutants (2B4, *sul1*Δ clone 1 and clone 2) was determine by two-way ANOVA, * p < 0.05, ** p < 0.01, *** p < 0.001, **** p < 0.0001. The highest p-value within all 3 comparisons is indicated. In K, differences in the abundance of detected indoles between WT and *sul1*Δ grown in the presence of tryptophan were determined using Student’s t-test with FDR correction, * FDR < 0.05. **See also Supplementary Figure S5.**

To confirm the phenotype of the *sul1* random insertional mutant, a *sul1*Δ targeted-deletion mutant was then generated in *M. furfur* CBS14141 with *Agrobacterium*-mediated transformation by replacing the entire *SUL1* open reading frame (**Figure 5C**, bottom). Consistent with the results of the genetic screen, both the *sul1* insertional mutant 2B4 and two independent clones of the *sul1*Δ targeted mutant (*sul1*Δ#1, *sul1*Δ#2) displayed clear inability to produce brown pigments when tryptophan was the sole nitrogen source, both in liquid and solid mDix-mp and MM media (**Figure 5D-F**).

Because Sul1 is a predicted sulphate transporter, we next tested the ability of the *sul1* mutants to utilize different inorganic and organic sulphate sources in presence or absence of a nitrogen source (i.e. ammonium nitrate). Phenotypic analysis revealed that in comparison to the parental WT, *sul1* mutants displayed slightly reduced growth and smaller colonies on MM, and on MM supplemented with sodium sulfide (Na_2_S), magnesium sulfate (MgSO_4_), sodium sulfate (NaSO_4_) and potassium sulfate (KSO_4_), and with the organic sulfur sources methionine (Met) and cysteine (Cys) (**Figure 5G**, top). Conversely, the mutants displayed the same phenotype as WT on potassium metabisulfite (K_2_S_2_O_5_) and sodium sulfite (NaSO_3_), indicating that *M. furfur* Sul1 is a transporter of sulphide, sulphate and of the two sulfur-containing proteinogenic amino acids, but not of sulfur dioxide and sulfite. The presence of a nitrogen source did not impact the *sul1* mutant phenotypes (**Figure 5G**, bottom).

To correlate the observed phenotypes wit the pigmentation ability of *M. furfur*, we tested the impact of the distinct sulfur sources on pigment production by growing WT *M. furfur* in MM supplemented with tryptophan and replacing MgSO_4_ by MgCl_2_. As expected, the sulfur sources that affected growth of the *sul1* mutants (**Figure 5H-I**) also directly influenced pigment production in WT *M. furfur*. Sodium sulfide and magnesium sulfate led to stronger pigmentation compared to the no-sulphur control (MgCl_2_) and potassium bisulphite. Sodium sulfite and sulfide led to intermediate pigmentation, which appeared brighter and more orange-like compared to the magnesium sulphate condition **(Figure 5H**). Moreover, all tested sulphate sources significantly increased brown pigmentation compared to the MM medium without sulphate (**Figure 5I**). In contrast, the organic sulphate sources methionine and cycteine had no impact on pigmentation (**Supplementary** Figure 5A). Together, these findings indicate that the Sul1 transporter in *M. furfur* is critical for pigmentation by facilitating the uptake of inorganic sulfate and sulfide that probably serve as cofactor of the enzymes involved in AhR-agonists biosynthesis.

To gain deeper insights into how Sul1 affects indole production, we conducted a metabolomic analysis of the secretome from the *sul1*Δ#1 deletion mutant and the parental *M. furfur* CBS14141 after 4 days of growth in mDixon-mp supplemented with or without tryptophan. Principal component analysis showed clustering among the different samples (**Figure 5J**). The comparison of *sul1*Δ +Trp versus WT +Trp revealed 397 differential molecular features, with 142 upregulated and 255 downregulated (**Supplementary Table S2**), while *sul1*Δ +Trp versus *sul1*Δ without Trp resulted in 162 significant molecular features, with 79 upregulated and 83 downregulated (**Supplementary Table S3**). Among the detected indolic metabolites, malasseziazole C, indole-3-acetic acid, malassezin, D-indole-3-lactic acid, and malasseziacitrin were found exclusively in WT +Trp, while pityriazepin/pityriacitrin and malassezindole A were only detected in the *sul1*Δ mutant (**Figure 5K**). Malassezindole B, pityriazole and indole-3-carbaldehyde were present in both the *sul1*Δ mutant and WT +Trp, though malassezindole B significantly upregulated in the mutant, indole-3-carbaldehyde significantly downregulated and pityriazole not significantly different (**Figure 5K**). Notably, no tryptophan-derived metabolites were detected in either *sul1*Δ or WT without tryptophan (**Figure 5K, Supplementary Table S4**). Additionally, we detected a notable accumulation of tryptophan in the sul1Δ +Trp mutant, whereas tryptophan levels were barely not detectable in the WT +Trp (**Figure 5L**), suggesting that Sul1 catalyzes tryptophan utilization. While this does not indicate that Sul1 directly transports tryptophan, its absence may impair metabolic pathways that normally consume or process tryptophan, leading to its accumulation. In conclusion, the majority of the tryptophan derived set of indoles produced by WT *M. furfur* are Sul1-dependent, as their absence in the *sul1*Δ mutant suggests that Sul1-mediated sulfate uptake is essential for their biosynthesis, while the limited indole production in the mutant, including alternative indoles not detected in WT *M. furfur*, may reflect a compensatory metabolic shift.

### Restoration of barrier integrity in inflamed skin by *M. furfur* depends on indole production by the fungus

To confirm the role of indole production by *M. furfur* in the skin barrier restoration, we first assessed the consequences of Sul1 deficiency in *M. furfur* on AhR signaling. Both clones of the *M. furfur sul1*Δ mutant exhibited a strong defect in activating AhR signaling in HaCaT keratinocytes in comparison to WT *M. furfur*, which induced strong *Cyp1a1* expression when grown in the presence of tryptophan (**Figure 6A**). The defect was even more pronounced when comparing the response to soluble tryptophan metabolites produced by *sul1*-deficient vs. Sul1-sufficient *M. furfur* (**Figure 6B**). Reduced AhR activation by secreted metabolites from *M. furfur sul1*Δ compared to those from WT *M. furfur* was further confirmed by means of the GFP reporter cell line (**Figure 6C**).

**Figure 6.**
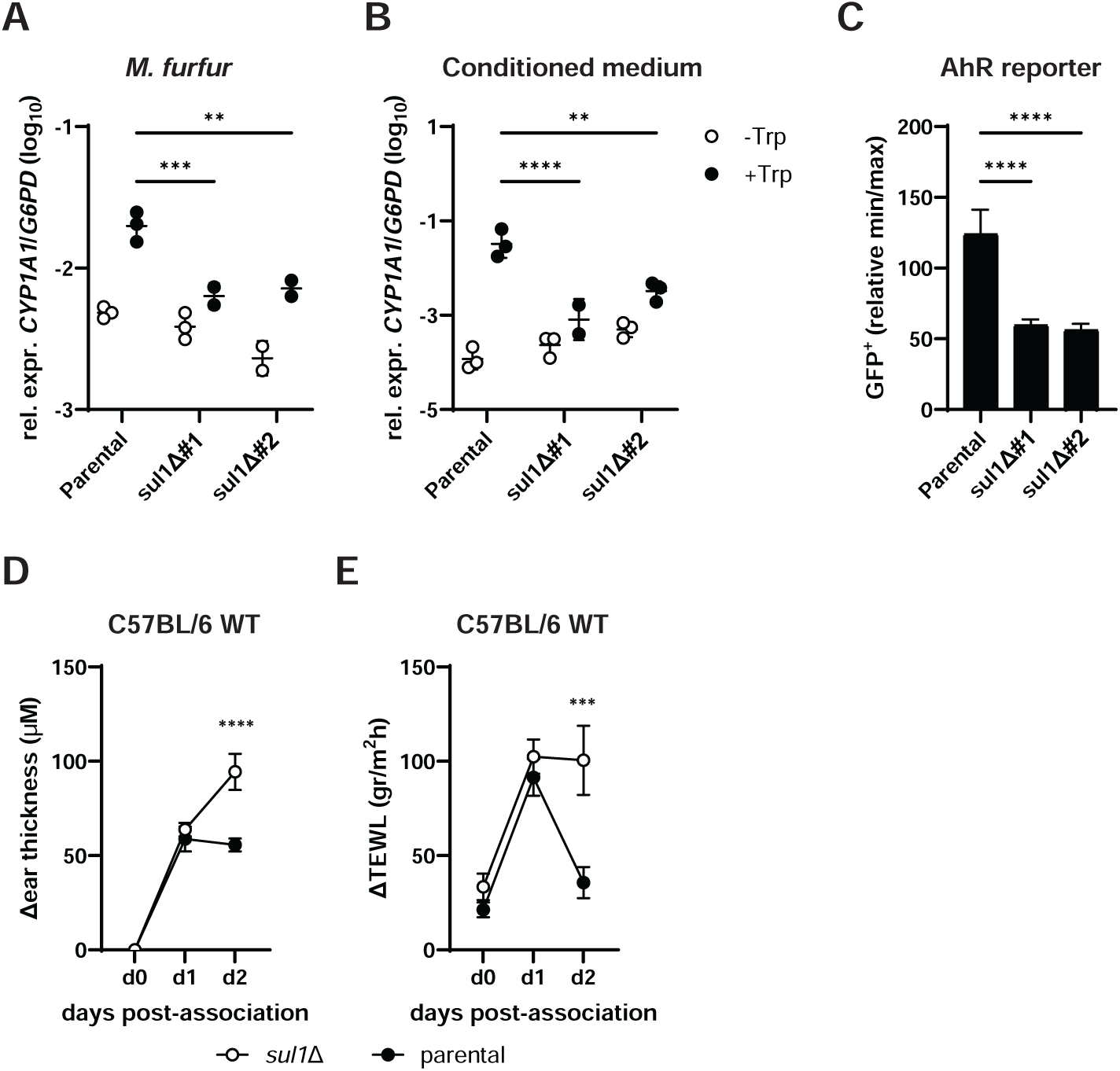
Restoration of barrier integrity in inflamed skin by *M. furfur* depends on indole production by the fungus. **A. – B.** *CYP1A1* expression by HaCaT keratinocytes after 24 hours of infection with *M. furfur sul1*Δ mutant clone 1 and clone 2 and the WT parental strain CBS14141 (A) or exposure to conditioned medium from the same strains (B) grown in presence or absence of tryptophan. n = 3 / group. The mean of each group is indicated. **C.** GFP expression by the reporter cell line H1G1-1C3 in response to 24 hours of stimulation with conditioned medium from *M. furfur sul1*Δ mutant clone 1 and clone 2 or the WT parental strain grown in presence of tryptophan. Expression levels are displayed relative to unstimulated and FICZ-stimulated cells. n = 8 / group. The mean + SD is indicated. **D. – E.** The ear skin of WT C57BL/6 mice was mildly tape stripped prior to epicutaneous administration of *M. furfur sul1*Δ mutant clone 1 and the WT parental strain pre-grown in presence of tryptophan. Increase in skin thickness (D), and TEWL (E). n = 8 / group pooled from 2 independent experiments (D, E). In A – C, the statistical significance of differences between mutant clones and WT was determined by two-way ANOVA (A, B) or One-way ANOVA (C). In D – E, the statistical significance of differences between the *M. furfur* +Trp and *M. furfur* -Trp groups was determined by two-way ANOVA. * p < 0.05, ** p < 0.01, **** p < 0.0001.

We then turned back to our *in vivo* skin inflammation model to assess the functional consequences of impaired AhR activation in response to *sul1*Δ *M. furfur*. Unlike WT *M. furfur*, which curbed skin inflammation (reduced skin thickness) and restored barrier integrity (reduced TEWL) in tape-stripped skin, when grown in the presence of tryptophan, the *sul1*Δ mutant failed to do so as indicated by increased skin thickness and elevated TEWL (**Figure 6, D - E**). Together, these results confirmed that indole production by *M. furfur* is essential for AhR activation in keratinocytes and for the AhR-mediated skin-protective effects of *M. furfur*.

## DISCUSSION

As a member of the skin microbiota, *Malassezia* stands out by its abundances. Unlike any other human colonizer, it largely dominates the skin fungal community [26]. *Malassezia* has uniquely adapted to the conditions of mammalian skin environment and thrives on host-derived lipids. Its persistence across different skin body sites suggests that it may play a role in skin physiology, yet its contributions to specific structural and immunological barrier functions have remained unexplored. Here, we demonstrate that *M. furfur*, via the secretion of indole metabolites, supports epidermal barrier homeostasis. This represents the first demonstration of a mutualistic role of *Malassezia*, with a direct effect on the host. These findings both challenge and complement the prevailing view that *Malassezia* is an opportunistic pathogen, frequently implicated in skin disorders that range from mild conditions such as dandruff to chronic inflammatory diseases like sebhorreic and atopic dermatitis (e.g.,[56]). Understanding the dual role of *Malassezia* in health and disease provides a more nuanced understanding of *Malassezia*–host interactions and emphasizes the importance of microbial-derived metabolites in maintaining skin health.

Although the detailed mechanisms of pathogenesis underlying *Malassezia*-associated skin diseases remain to be uncovered, the fungus is thought to contribute to disease progression by secreting proteases, lipases, and phospholipases, which are found selectively upregulated in diseased skin [57, 58]. These enzymes can exacerbate barrier defects and amplify inflammation, key features of diverse skin disorders [59–63]. In atopic dermatitis, sensitization to *Malassezia* with fungal allergens driving IgE and Th2-mediated inflammation [64] is a further hallmark of disease emphasizing the relevance of *Malassezia* in disease.

By identifying a fungal mechanism through which *Malassezia* restores skin barrier integrity, our study provides a critical new perspective on the impact of this prevalent skin colonizer for the host. We uncover a new biochemical pathway in *Malassezia* that controls the production of indole metabolites and by means of gene-deletion mutants in both, the fungus and the host, demonstrate that these compounds actively influence host signaling.

Among the common *Malassezia* species in human skin, *M. furfur* is not the most prevalent one. However, its relative abundance increases in atopic and seborrheic dermatitis [65, 66], and isolates from diseased skin have been reported to exhibit enhanced indole production [22, 24]. *M. furfur* strains obtained from lesional skin of SD patients produced higher levels of indoles in response to tryptophan in comparison to isolates from healthy individuals [22]. This led to the assumption that indole production by *Malassezia* is a pathogenicity trait, contributing to disease progression [22, 67, 68]. Our findings, however, challenge this hypothesis and instead support a host-protective role for fungal indoles. Rather than driving pathogenesis, indole production may be an adaptive mechanism of the fungus that gets activated under pathological conditions to restore tissue homeostasis. This perspective is strongly supported by our results showing that *Malassezia*-derived indoles induce host signaling pathways, such as AhR, to enhance the skin barrier function and dampening inflammation. Such a mechanism suggests that, under conditions of barrier disruption and immune dysregulation, *M. furfur* may contribute to epidermal resilience rather than exacerbate disease, thereby positioning its metabolic activity as a compensatory response to environmental or host-derived stressors.

The AhR-mediated effects of *M. furfur*-derived indoles on the skin barrier align with previous findings demonstrating that microbial and dietary indoles contribute to barrier integrity in both, skin and gut [14, 53, 69]. However, in most of these studies, the microbial species responsible for indole production remained unidentified, leaving the specific microbes contributing to AhR activation largely speculative. In contrast, our study identifies *M. furfur* as a defined fungal source of indoles and directly links its metabolic activity to AhR-dependent skin barrier regulation. To confirm the relevance of *M. furfur*’s tryptophan metabolism in this process, we generated indole-deficient fungal mutants. The lack of barrier restoring effects in the mutants confirmed that fungal-derived indoles actively contribute to AhR-mediated skin barrier homeostasis, rather than being incidental metabolic byproducts.

*Malassezia*’s ability to produce indoles is regulated not only by tryptophan availability but also by that of sulfur. This suggests that *Malassezia*’s indolic metabolism is highly adaptable and responds to specific environmental cues present in the skin. Sulfur-containing compounds are available in the skin through keratin degradation, sebum components, and sweat-derived thiols [70, 71]. Interestingly, while tryptophan metabolism is essential for indole biosynthesis across multiple fungal species, key enzymes previously implicated in indole production in other fungi, such as the aminotransferase *TAM1/ARO8/AroH/* mediating the first step of tryptophan degradation in *U. maydis, C. glabrata* and *Aspergillus fumigatus*, respectively, [72–74], appear to be dispensable or reduntant with other aminotransferases in *M. furfur*, despite a biochemical confirmation on *M. furfur* CBS 1878 [75]. Instead, we identified a sulfate-dependent pathway as a crucial determinant of indole biosynthesis in *M. furfur*, a finding that corroborates a previous findings in *Ustilago maydis*, in which a forward genetic screen identified a mutant with impaired pigmentation with mutation in a gene related to sulfur metabolism [74]. Therefore, similarities in tryptophan catabolism are shared between *M. furfur* and *U. maydis*, a basidiomycete plant pathogen taxonomically related to *Malassezia*. In contrast, *Cryptococcus neoformans*, a basidiomycete related to *Malassezia* and *Ustilago*, relies on a laccase-dependent pathway for tryptophan degradation [77]. These data suggest that fungal tryptophan degradation pathways are fundamental, with their evolution being shaped by environmental factors that drive fungal adaptation. Given the abundance of sulfur-containing compounds in the skin, it is plausible that the integration of sulfur metabolism into *Malassezia*’s tryptophan metabolic pathway represents an adaptation to its host-associated lifestyle, paralleling the evolutionary adaptation of *Ustilago*, which thrives in sulfur-rich plant environments. The dependence of *Malassezia* indole production on sulfur suggests that host-derived sulfur compounds may modulate the yeasts ability to generate AhR ligands, raising important questions about whether sulfur availability may modulate *Malassezia*’s role in skin homeostasis and inflammation.

Given the anti-inflammatory consequences of AhR signaling in barrier tissues [78, 79], AhR ligands are increasingly being explored as pharmacological agents [80]. Tapinarof, a topical AhR agonist approved for the treatment of plaque psoriasis [81] is currently undergoing clinical trials for moderate to severe atopic dermatitis [82, 83]. Despite promising therapeutic potential, however, targeting AhR in inflammatory disorders is not a guaranteed success. The pleiotropic nature of AhR ligands means that their effects can vary dramatically depending on the tissue, cell type, and host condition, and also include pro-inflammatory consequences [84–87]. Nevertheless, most examples of AhR stimulation in the skin are associated with host-beneficial effects, particularly by enhancing barrier integrity and tissue homeostasis.

Together, our findings reveal a dynamic interplay between microbial signals and keratinocyte-specific AhR activation in shaping the skin’s barrier properties. The ability of microbial-derived metabolites, particularly fungal indoles, to modulate epidermal differentiation, immune responses, and barrier restoration underscores a previously underappreciated role for the skin mycobiota in maintaining homeostasis. Expanding the scope of microbiota research to include fungal-derived bioactive compounds will be critical for understanding their contributions to skin physiology and developing microbiome-targeted therapies to enhance skin health and restore barrier integrity in common inflammatory cutaneous disorders.

## STAR METHODS

### Experimental model and study participant details

#### Ethics statement

All mouse experiments in this study were conducted in strict accordance with the guidelines of the Swiss Animals Protection Law and were performed under the protocols approved by the Veterinary office of the Canton Zurich, Switzerland (license number ZH142/2021). All efforts were made to minimize suffering and ensure the highest ethical and humane standards according to the 3R principles [88]. Foreskin biopsies were obtained from males, collected with informed written consent upon approval from local ethical committees (Kantonale Ethikkommission Zürich, approval numbers: 2015-0198 and 2024-01030) and were conducted according to the Declaration of Helsinki Principles.

#### Animals

Female WT C57BL/6j female mice were purchased from Janvier Elevage and used for experiments at 6-14 weeks for age. *AhR*^−/−^ mice were generated by crossing *AhR^fl/fl^*(Jax #006203) [89] and CMV-cre (Jax #006054) [90]. *AhR*^Δ*K14*^ mice were generated by crossing *AhR^fl/fl^* (Jax #006203) and K14-cre [91]. Female and male (conditional) knockout mice were used at 6-14 weeks of age for experiments and littermates were used as controls. All mice were kept in individually ventilated cages under specific pathogen-free conditions at the Institute of Laboratory Animals Science (LASC, University of Zurich). Animals were allowed to acclimatize for 1 week after arrival in the BSL2 animal experimentation unit of LASC before starting experiments. Only animals in good health were included in experiments. Experiments with WT mice used a randomized design; experiments with genetically modified mice were not done fully randomized due to variable distribution of the different genotypes in each litter. Colonized and non-colonized animals were kept separately to avoid cross-contamination. Sample size was chosen by the Fermi’s approximation and based on experience.

#### Fungal strains

*M. furfur* strain JPLK23 (CBS 14141) and *M. sympodialis* strain ATCC 42132 were originally obtained from Joseph Heitman (Duke University Medical Center, Durham, NC). *M. pachydermatis* strain ATCC 14522 (CBS 1879) and *M. globosa* MYA-4612 (CBS7966) were purchased from ATCC. *M. restricta* CBS 7877 was obtained from Philipp Bosshard (University Hospital Zurich, Switzerland). All strains were maintained at 30 °C, 180 RPM in liquid modified Dixon medium (mDixon) containing 36 gr Malt Extract (Sigma Aldrich), 20 g desiccated Ox-bile (Sigma Aldrich), 10 ml Tween-40 (Sigma Aldrich), 6 g peptone (Oxoid), 2 ml Glycerol (Honeywell), 2 ml oleic acid (Sigma Aldrich) per 1 liter or in mDix-mp, before culturing in assay-specific media, as specified in the Methods details below and in results section. To quantify fungal density, cultures were washed twice and resuspended in PBS. Number of yeast cells were determined by measuring OD_600_ (1 OD_600_ = 5 x 10^6^ yeast cells / ml).

#### Cell lines and maintenance

The human keratinocyte cell lines HaCaT [92], N/TERT1 [93], and HPK were used for infection with *Malassezia* and treatment with *Malassezia*-derived metabolites. The immortalized hepatocyte cell line H1G1-1C3 [41], kindly provided by Professor Michael S. Denison, was used for AhR reporter assays. 3T3-J2 feeder cells were used to prepare HPK. HPKs were prepared from foreskin biopsies. Experiments were conducted on cells at passage 4 for HPK, between passages 30 and 50 for HaCaT, passages 38 to 45 for N/TERT1, and passages 10 to 30 for H1G1-1C3. For routine culture, HaCaT and H1G1-1C3 cells were maintained in DMEM high glucose supplemented with 10% FCS and 1% Pen/Strep, while N/TERT1 and HPK cells were cultured in K-SFM (Invitrogen) supplemented with bovine pituitary extract (half-tube supplied with K-SFM), epidermal growth factor (0.2 ng/mL), and calcium chloride (0.3 mM), without antibiotics or antimycotics. All cells were incubated at 37°C, 5% CO₂ in a humid chamber. For routine passaging, cells were washed with PBS and, detached with 0.05% Trypsin-EDTA. The trypsin reaction was stopped with DMEM containing 10% FCS. N/TERT1 cells were also used for 3D human epidermal equivalents (HEEs).

### Methods details

#### Pigments production by *M. furfur*

To induce pigmentation and indole production, *Malassezia* strains were grown in mDix-mp (for 1 L, 36 gr Malt Extract, 10 g desiccated Ox-bile, 10 ml Tween-40, and 2 ml Glycerol), MM (for 1 L, 15 g Glucose, 1 g KH_2_PO_4_, 0.5 g KCl, 0.5 g MgSO_4_, 4 mL Tween 60, 1 mL Tween 20, 4 g Ox-bile) or Tween 80 agar (30 ml of Tween 80 for 1 L) supplemented with 5 mg/ml of L-tryptophan, if not specified otherwise. For solid media, 2% agar was used.

Grown and pigment production experiments were carried out in MM with 15 g/L of glucose. On a daily basis, 250 µL of each cellular suspension were withdrawn and 100 µL used to monitor the growth at OD_600_, and 100 µL were centrifuged to collect cell-free supernatant used to monitor pigmentation at OD_400_; the ratio OD_400_/OD_600_ is used to express pigments production on the growth.

Pigmentation of the WT strain CBS14141 in the presence of different sulfur sources was carried out in MM with 15 g/L of glucose and in which MgSO_4_ was replaced by MgCl_2_, with and without ammonium nitrate (20 mM), and with 2 mM of the following sulfur sources: sodium sulfide (Na2S), potassium metabisulfite (K2S2O5), sodium sulfite (Na2SO3), magnesium sulphate (MgSO4 x 7 H20 and MgSO4 x 1H20), sodium sulphate (Na2SO4), and potassium sulphate (KSO4); the aminoacids methionine and cysteine were used at 5 mM.

#### Epicutaneous colonization of murine ear skin

Mice were anesthetized by injection of 65 mg/kg Ketamin and 13 mg/kg Xylazin and the dorsal side of their ears was subjected to mild tape stripping using Transpore™ Hypoallergenic tape (3M; 5 rounds per ear) to disrupt the stratum corneum. Prior to colonization, ear thickness and transepidermal water loss (TEWL) were measured using an Oditest S0247 0–5 mm measurement device (KROEPLIN) and a Tewameter® TM Nano device (Courage + Khazaka), respectively, to establish baseline values. A 100 µL yeast suspension (corresponding to 10LJ yeast cells) was then applied epicutaneously to the dorsal side of each ear. Control groups received olive oil without *M. furfur*. Ear thickness and TEWL were measured prior colonization and then daily throughout the experiment. Ear thickness was monitored using the Oditest S0247 0-5 mm measurement device (KROEPLIN). TEWL was measured with a non-invasive probe (Tewameter® TM Nano, Courage+Khazaka, Cologne, Germany) by placing the probe in the same place of the dorsal side of the ear. Mice were anesthetized with vaporized isoflurane during measurements. For determining the skin fungal loads, the ear tissue was homogenzied in water supplemented with 0.05 % Nonidet P40 (AxonLab), using a Tissue Lyser (Qiagen), and plated on mDixon agar for incubation at 30°C for 3 to 4 days.

#### Epidermal sheets preparation and stimulation

Mouse ears were collected and the dorsal side was mechanically separated from the ventral side. The separated tissue was then enzymatically digested using 3 U/ml Dispase II (Roche) diluted in PBS at 37°C for 1 hour before carefully separating the epidermis from the dermis. Epidermal sheets were incubated in conditioned medium for 24 hours at 37°C and 5% CO_2_. After incubation, epidermal sheets were collected, snap-frozen in liquid nitrogen and stored at -80°C until RNA isolation was performed.

EROD assay

CYP1A1 enzymatic activity was measured using the EROD assay, as previously described [94], with modifications. Briefly, epidermal sheets were placed in a 96-well plate containing 100 µL of 2 µM 7-ethoxyresorufin (7-ERO, Sigma-Aldrich) in sodium phosphate buffer (50 mM, pH 8.0) per well. The plate was incubated at 37 °C for 20 minutes and the reaction was terminated by adding fluorescamine (150 µg/mL in acetonitrile, Sigma-Aldrich). Resorufin formation was quantified using a resorufin (Sigma-Aldrich) standard curve on a plate reader (Tecan) with excitation and emission wavelengths of 535 and 590 nm, respectively. After termination of the assay, epidermal sheets were snap-frozen in liquid nitrogen and stored at -80 °C for RNA isolation.

#### Neutrophil quantification

Ears were collected from colonized mice three days post fungal association, cut into small pieces, and enzymatically digested in Ca²⁺- and Mg²⁺-free Hank’s medium (Life Technologies) supplemented with Liberase TM (0.15 mg/mL, Roche) and DNase I (0.12 mg/mL, Sigma-Aldrich) for 1 hour at 37°C. Single-cell suspensions were obtained by mechanical dissociation using a 70 µm cell strainer (Falcon). For neutrophil identification, cells were stained with AF700 Anti-CD45.2 (clone 104), Pacific Blue anti-Ly6G (clone 1A8), and PE-Cy5 anti-CD3ε (clone 145-2C11) at 4°C for 30 minutes in the dark. LIVE/DEAD

Fixable Near-IR stain (Life Technologies) was used to exclude dead cells. Flow cytometry was performed using a CytoFLEX (Beckman), and data were analyzed with FlowJo software. Gating strategies followed the guidelines for the use of flow cytometry and cell sorting in immunological studies [95], including pre-gating on viable and single cells (**Supplementary Figure S3E**).

#### Histology

Mouse tissues were fixed in 4% PBS-buffered paraformaldehyde overnight and subsequently embedded in paraffin and cut. Sagittal sections (9 µm) were then stained with hematoxylin and eosin (H&E) reagent, followed by counterstaining with hematoxylin. Slides were mounted using Pertex mounting medium (Biosystem) according to standard protocols. Slides were mounted in an aqueous mounting medium according to standard protocols. All histological images were acquired using a digital slide scanner (NanoZoomer 2.0-HT, Hamamatsu) and analyzed with NDP.view2 software. Epidermal thickness was measured by randomly selecting five areas per image, each separated by 1 µm along the epithelium, and measuring the thickness of the epidermis at these position with the ruler tool of the NDP.view 2 software.

#### *Malassezia* mutant generation

To generate random insertional mutants of *Malassezia furfur*, *Agrobacterium tumefaciens*-mediated transformation was performed using the binary plasmid pAIM2, which carries the nourseothricin (NAT) resistance marker, following a previously described [38]. Briefly, *M. furfur* CBS 14141 was cultured at 30°C for two days, while *A. tumefaciens* was grown overnight in Luria-Bertani (LB) medium supplemented with 50 µg/mL kanamycin. Both *M. furfur* and *A. tumefaciens* cultures were subsequently diluted to an OD_600_ of 1 and mixed at *Malassezia:Agrobacterium* ratios of 2:1, 5:1, and 10:1. The mixtures were pelleted at 4000 rpm for 15 min, resuspended, and spotted onto nylon membranes placed on modified IM (mIM) plates containing 100 µM acetosyringone, 0.4% v/v Tween 60, and 0.1% v/v Tween 20. After three days at 24°C, cells were recovered, washed, and spread onto mDixon with NAT (100 µg/mL), ampicillin (100 µg/mL), and cefotaxime (300 µg/mL). NAT-resistant transformants were arrayed in 96-well plates and replica-plated onto mDixon-mp with 0.5 mg/mL L-tryptophan. Mutants with impaired pigmentation were identified through visual inspection and retested under identical conditions. T-DNA insertion sites were determined by inverse PCR [96] using genomic DNA extracted from overnight liquid mDixon cultures with a CTAB-based method [97]. Approximately 2 µg of DNA was digested with the restriction enzymes ApaI, EcoRI, XhoI (New England Biolabs), self-ligated with T4 DNA Ligase (New England Biolabs) at 4 °C overnight, and amplified using primers ai76-ai77 or ai076-M13F/ai077-M13R **(Key Resource Table**) for Sanger sequencing. BLAST analysis against the *M. furfur* CBS 14141 genome (GCA_023510035.1) to identify T-DNA insertion sites (Theelen et al., 2022) was performed. The hit gene was subjected to blast searches on GenBank and Saccharomyces Genome Database to infer its predicted function; *Saccharomyces cerevisiae* gene nomenclature was used.

#### Targeted mutagenesis of the *SUL1* gene

Plasmid for targeted mutagenesis of *M. furfur SUL1* through *A. tumefaciens*-mediated transformation were assembled in *S. cerevisiae* using the binary vector pGI3 as previously reported [37]. The *NAT* cassette was amplified from plasmid pAIM1 with primers JOHE43277 and JOHE43278, and the 5LJ and 3LJ flanking regions for homologous recombination were amplified from *M. furfur* CBS14141 genomic DNA with primer pairs PVCB164-PVCB165 and PVCB166-PVCB167, respectively. The PCR products and the digested (KpnI and BamHI) binary vector pGI3 were assembled into S*. cerevisiae*, and resulting transformants subjected to PCR with primers JOHE43279–JOHE43281 and JOHE43280–JOHE43282 to assess correct recombination at the 5LJ and 3LJ regions, respectively. Positive clones of *S. cerevisiae* were grown ON in YPD and subjected to phenol-chloroform-isoamyl alcohol (25:24:1) plasmid extraction [98]. The plasmid DNA, named pGI82, was then introduced into the *A. tumefaciens* EHA105 strain by electroporation, and transformants were selected on LB + 50 µg/mL kanamycin. *A. tumefaciens-*mediated transformation was performed as described above. *M. furfur* transformants resistant to NAT were subjected to molecular characterization to assess correct replacement of the target loci. Diagnostic PCRs were carried out with primers JOHE45213 or JOHE45874 in combination with specific primers for the *NAT* gene (JOHE43281 and JOHE43282, respectively), and with primers JOHE45215-JOHE45216 specific for the internal region of *CDC55.* Two independents *sul1*Δ mutants were selected and used for phenotypic evaluation together with the insertional *sul1* mutant (strain 2B4) and the parental CBS 14141. Pigments production was assessed in liquid and solid mDix-mp and MM as described above. The ability to growth on different sulphur sources was assessed by spotting 3 µL of 1:10 dilutions of each cellular suspension on MM medium (note that MgSO4 was replaced with MgCl2) with glucose with and without ammonium nitrate in the presence of sulfur sources at 2 mM of sodium sulfide (Na2S), potassium metabisulfite (K2S2O5), sodium sulfite (Na2SO3), magnesium sulphate (MgSO4 x 7 H20, and MgSO4 x 1H20), sodium sulphate (Na2SO4), and potassium sulphate (KSO4); methionine and cysteine were used at 5 mM. Plates were incubated at 30°C.

#### Preparation of conditioned medium

*Malassezia* strains were cultured in liquid mDixon medium for 2 days at 30°C with agitation at 180 RPM. 2.5 × 10LJ yeast cells were then inoculated onto Tween 80 agar plates containing or not-containing 5 mg/ml L-tryptophan and incubated at 30°C for 5 days. Fungal cells were collected using a cell scraper and resuspended in 4 mL of F12 medium (Gibco) supplemented with 1% FCS and 1% Pen/Strep (complete F12). For HEE experiments, yeast cells were resuspended in CnT Prime 3D Barrier medium (CELLnTEC). To remove fungal cells, the samples were passed through a 0.22 µm filter, and sterility was verified by plating on mDixon agar.

#### Human epidermal equivalents

For the generation of human epidermal equivalents (HEEs), cryopreserved N/TERT1 aliquots containing 1×10^6^ cells at passage numbers between 38 and 41 were freshly thawed. Cells were grown in K-SFM until confluency. Cells were detached, collected and counted. The cell concentration was adjusted to 2×10^6^ cells/ml in CnT Prime medium (CELLnTEC). 24-well cell culture plates were prepared by hanging empty cell culture inserts (ThinCertTM for 24 Well Multiwell Plates, Greiner Bio One) into wells containing 1 ml of CnT Prime medium. 100 µl of cell suspension was seeded into each cell culture insert. After incubation at 37°C, 5% CO_2_ for 24h to 48h the cells were checked for confluency. When 100% confluency was reached, the CnT Prime in the bottom well was replaced by 1 ml of 3D barrier medium (3D barrier medium, CELLnTEC). The medium in the cell culture insert was cautiously aspirated while avoiding touching the cell layer and likewise replaced with 100 µl of 3D barrier medium. After incubation overnight at 37°C, 5% CO_2_, the medium from the cell culture insert was aspirated to start the air-liquid-interface. The cell culture plates were left in the incubator at 37°C, 5% CO_2_ and the medium in the bottom well was replaced with 1 ml of fresh differentiation medium 3 times per week until start of treatment.

#### Isolation and culture of HPK

HPK were prepared as described [99]. Briefly, the foreskin biopsy was disinfected by a short incubation in ethanol, 5 min incubation in PBS 10% antibiotic/antimycotic (A/A, Thermo Fisher Scientific), 5 min in PBS 1% A/A and finally washed in pure PBS. Fat was removed from the skin biopsy and the tissue cut into small pieces. The skin pieces were then incubated for 2h in DMEM containing 1% A/A (without FBS) and subsequently in 4 U/ml Dispase II (Roche) in PBS, overnight at 4 °C. Separation of dermis and epidermis was performed the next day, and the epidermis was incubated in a PBS solution containing 0.25% Trypsin/EDTA (Thermo Fisher Scientific) for 20 min at 37 °C. A single cell suspension of keratinocytes was obtained by pipetting the epidermis up and down in DMEM containing 25% FBS and 1% A/A and passing the cell suspension through a 100 μm cell strainer. Cells were centrifuged (170 x g, 3 min, RT) and resuspended in complete keratinocyte medium (3 parts DMEM, 1 part Ham’s F12 (Thermo Fischer Scientific), 10 % FBS, 1 % A/A, 20 μg/ml adenine (Sigma, Munich, Germany), 5 μg/ml apo-Transferrin (Sigma), 2 nM 3,3’,5-Triiodo-L-thyronine (Sigma), 200 ng/ml hydrochortison (Sigma), 100 pg/ml cholera toxin (Sigma), 5 μg/ml insulin (Sigma), 10 ng/ml EGF (Sigma)). Cells were counted and 10^6^ keratinocytes were plated on top of 2×10^5^ mitotically-inactivated J2 feeder cells. After amplification of HPKs for two passages with 3T3-J2 feeder cells, they were cultivated in K-SFM (Thermo Fisher Scientific) with EGF and BPE.

#### Cas9 generation of *AHR* KO N/TERT1

*AHR* KO in N/TERT1 cells were generated according to [100]. Briefly, the 20-nucleotide gRNA targeting sequence for *AHR* was designed using the free online software ChopChop (https://chopchop.cbu.uib.no/) and Off-Spotter (https://cm.jefferson.edu/Off-Spotter/). Single-guide RNA (sgRNA) with the designed *AHR* targeting sequence and a negative control sgRNA (TrueGuide sgRNA Negative Control, non-targeting 1 (Invitrogen)) were obtained from Invitrogen and prepared according to the manufacturer. For assembling the RNP complexes, sgRNA and 5 μg/μL TrueCut HiFi Cas9 Protein (Invitrogen) were mixed at a 1:1 molar ratio: 24 pmol sgRNA and 24 pmol Cas9 per 0.15 × 10^6^ cells in 10 μL. The RNP mixture was incubated at room temperature for 20 min before addition of 0.15 *×* 10^6^ cells in 5 μL buffer R, followed by electroporation. For electroporation, the Neon Transfection System (Invitrogen) and the Neon Transfection System 10 μL Kit (Invitrogen) were used applying 1700 Volts/20 ms pulse width/1 pulse. The electroporated cells were seeded in a T150 flask with 30 mL K-SFM complete and expanded without passaging for at least a week. Medium was changed every two to three days until experiments.

#### Keratinocytes and HEE infection and treatment with conditioned medium

For experiments using monolayer keratinocyte cell lines, 5 × 10LJ cells per well were seeded in 100 µL of medium in 96-well plates, using DMEM for HaCaT and HPK cells or K-SFM for N/TERT1 cells, and incubated overnight at 37°C with 5% CO₂. The following day, the medium was refreshed, and 2 × 10LJ *M. furfur* yeast cells, resuspended in F12 medium, were added on top of the keratinocytes. For human epidermal equivalents (HEEs), *M. furfur* was added at the same concentration, incubated for 30 minutes to allow the fungi to settle, and the medium was carefully removed using a micropipette. For cond. med. treatment, the culture medium was replaced with 100 µL of cond. med. in monolayer cultures, while for HEEs, the medium in the bottom well was replaced with 1 mL of c.m. FICZ (1 µM in F12 medium) was used as a positive control for AhR activation.

#### AhR reporter cell assay

For AhR activation, 5 × 10LJ H1G1-1C3 cells per well were seeded in 100 µL of medium in a 96-well plate and incubated overnight at 37°C with 5% CO₂. The following day, the medium was replaced with c.m., and FICZ at 1 µM in F12 was used as a positive control. Cells were incubated for 24 hours, and GFP fluorescence was measured using a plate reader (Tecan) with excitation at 485 nm and emission at 515 nm. The signal from untreated cells was subtracted from all values, and AhR activation was expressed as a percentage relative to the FICZ signal.

#### RNA isolation and RT-qPCR

Isolation of total RNA was done using TRI-Reagent (Sigma-Aldrich) according to the manufacturer’s instructions. For adherent cells, cells from four wells of a 96-well plate were pooled in 500 µl TRI-Reagent. Murine epidermal sheets and HEE samples were collected in 500 µl TRI-Reagent. Samples were homogenized using a TissueLyzer (Qiagen) for 6 minutes at 25 Hz, and RNA was isolated by phase separation using chloroform. Finally, RNA was precipitated using isopropanol, washed in 75 % EtOH, air-dried, and resuspended in ultra-pure H_2_O. cDNA was generated by RevertAid reverse transcriptase (ThermoFisher) and random nonamers. Quantitative PCR was performed using SYBR green (Roche) and a 7500 Real-Time PCR System (Applied Biosystems) instrument. The primers used for qPCR are listed in the **Key Resource Table**. All RT-qPCR assays were performed in duplicate, and the relative expression (rel. expr.) of each gene was determined after normalization to mouse *Actb* or human *G6CPD* housekeeping genes.

#### Conventional PCR

The excision of *Ahr*^fl/fl^ was assessed by multiplex PCR as previously described [89]. The reaction was performed using the forward primers OL4062 and OL4064, along with the reverse primer OL4088 (**Key Resource table**). The PCR products allowed differentiation between alleles, with the *Ahr*^fl/fl^ -excised allele (OL4062/OL4088) yielding a 180-bp band, the unexcised *Ahr*^fl/fl^ allele (OL4064/OL4088) producing a 140-bp band, and the WT allele generating a 106-bp band (OL4064/OL4088).

PCR amplification was carried out with an initial denaturation step at 95 °C for 3 min, followed by 35 cycles of denaturation at 95 °C for 15 sec, annealing at 65 °C for 15 sec, and extension at 72 °C for 15 sec. A final extension step was performed at 72 °C for 5 min.

#### Metabolomics

The *M. furfur* WT CBS14141 and the *sul1*Δ#1 mutant were grown in mDix-mp with and without 0.5 mg/mL of tryptophan for 4 days at 30 °C under shaking condition. Cultures were then centrifuged at 4000 rpm for 3 minutes, and the supernatants filter sterilized and maintained in ice until their use. Filter-sterilized *M. furfur* culture supernatants were extracted with ethyl acetate (EtOAc, GC–MS grade, Merck), followed by drying with anhydrous sodium sulfate (Na₂SO₄; Carlo Erba, Italy). The solvent was removed under vacuum at 37°C using a rotary evaporator (RV 10, IKA®-Werke GmbH & Co. KG, Staufen, Germany). Each extract was solubilized in methanol (MeOH, LC–MS grade, Fluka, Buchs, Switzerland) at a final concentration of 1 mg/mL and analyzed by LC–MS qTOF (Agilent Technologies). A volume of 7 µL was injected into an Agilent HP 1260 Infinity Series liquid chromatograph equipped with a diode array detector (DAD) and coupled to a quadrupole time-of-flight (qTOF) mass spectrometer. Analyses were performed according to Prigigallo et al. [101]. Raw data were pre-processed using MassHunter Profinder software 8.0 (Agilent Technologies), and mass spectra were aligned and log₁₀-normalized for intensity before statistical analysis. Samples were categorized based on strain and/or tryptophan supplementation and subjected to Principal Component Analysis (PCA). Significant metabolite accumulation was determined using Student’s t-test with false discovery rate correction (p < 0.05) and a fold change threshold of >2.0. Metabolite identification and annotation were performed by comparing monoisotopic data with a custom-built fungal database and previously published data. Three independent cultures of each strains were prepared, and each sample was injected two times.

### Statistical analysis

All statistical analyses were performed using GraphPad Prism version 10.4.1 for Windows (GraphPad Software, San Diego, CA, USA, www.graphpad.com) and built-in functions within the R statistical environment Rstudio version 2024.12.0. For metabolomics data, statistical analyses were done using Mass Profiler Professional (version 13.1.1, Agilent Technologies) and MetaboAnalyst version 6.0 (https://www.metaboanalyst.ca/). Data visualization was conducted using GraphPad Prism and ggplot2 [102] within the R statistical environment (Rstudio version 2024.12.0). Details of each analysis, including specific statistic analysis and p values, are provided in the figure legends.

### Key Resource Table

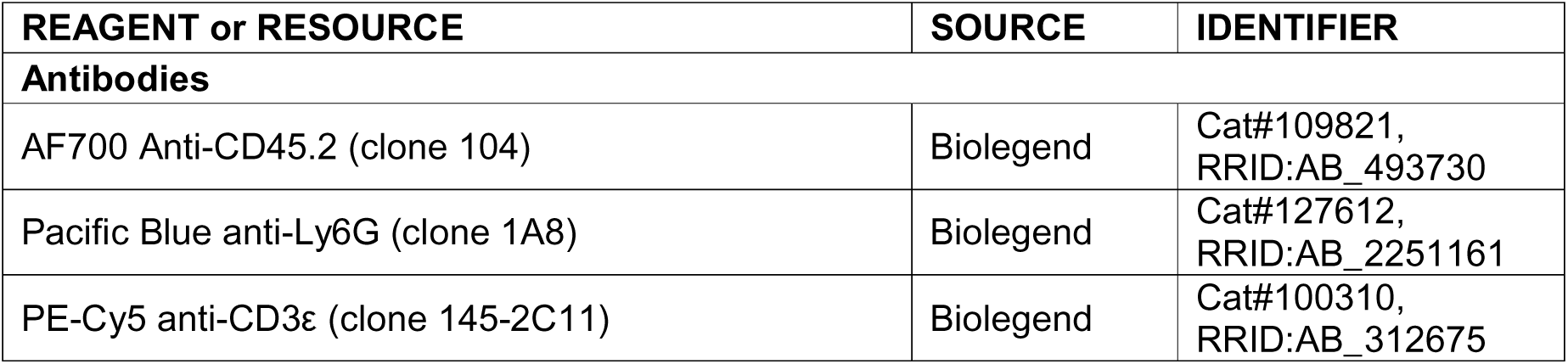
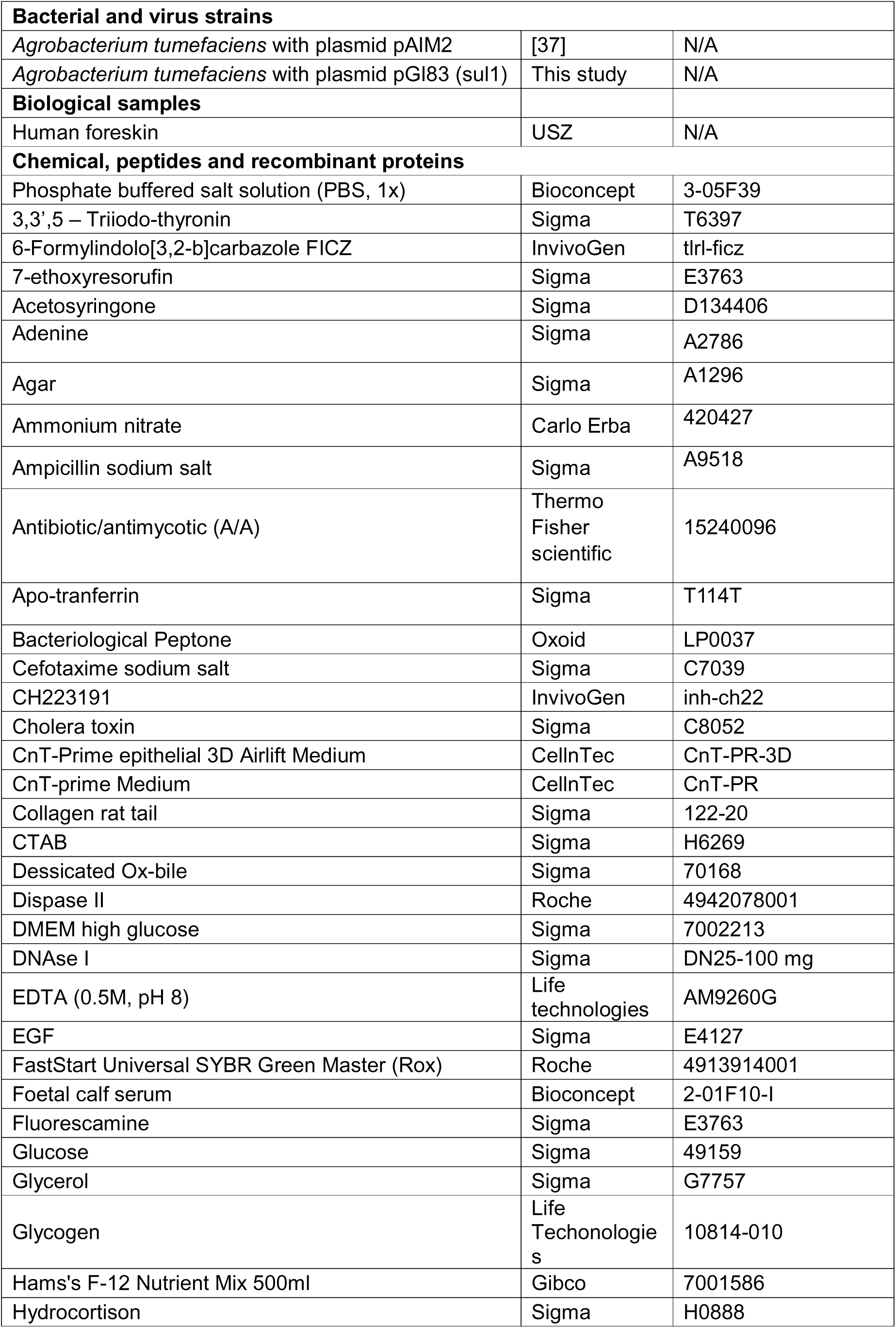
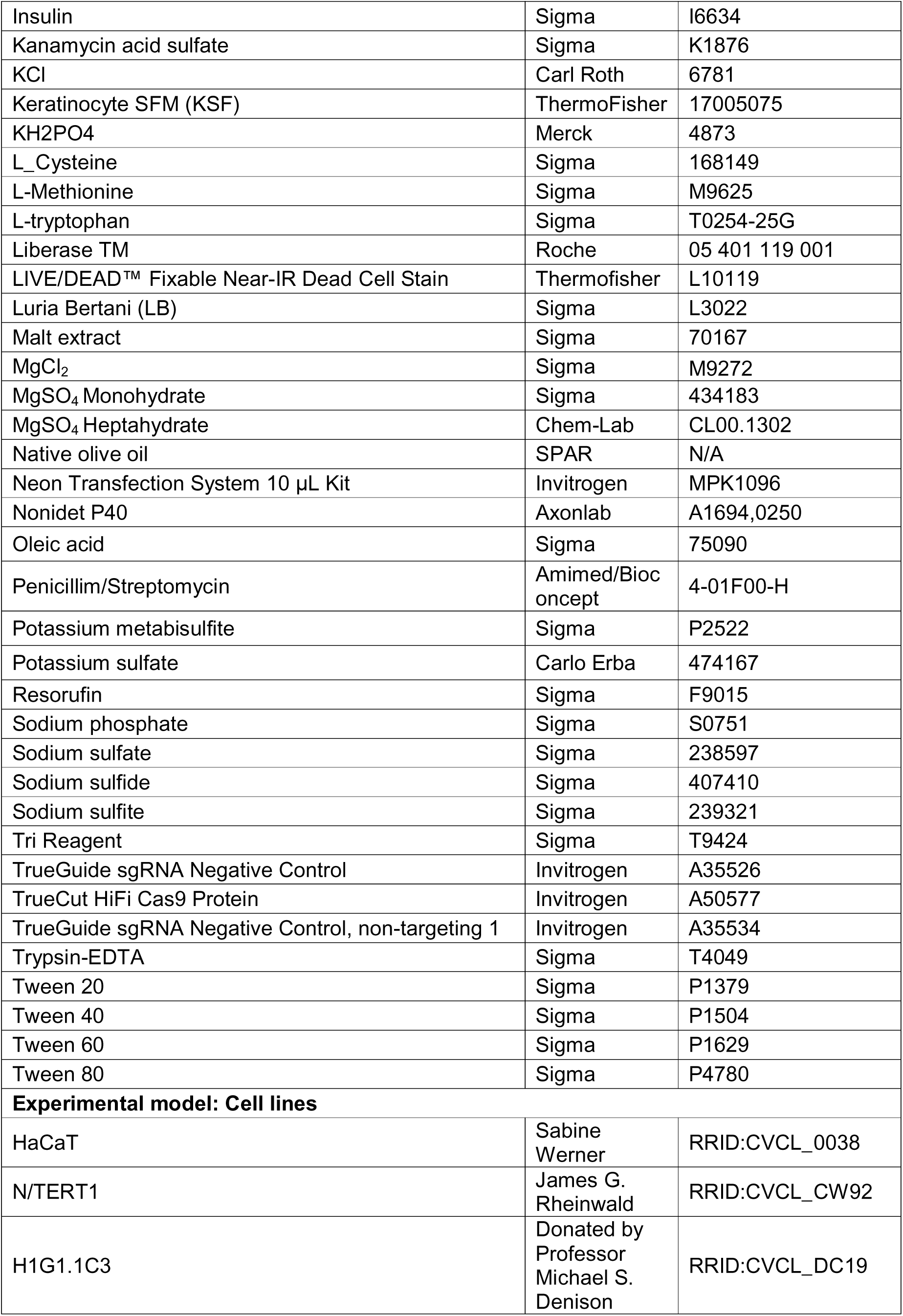
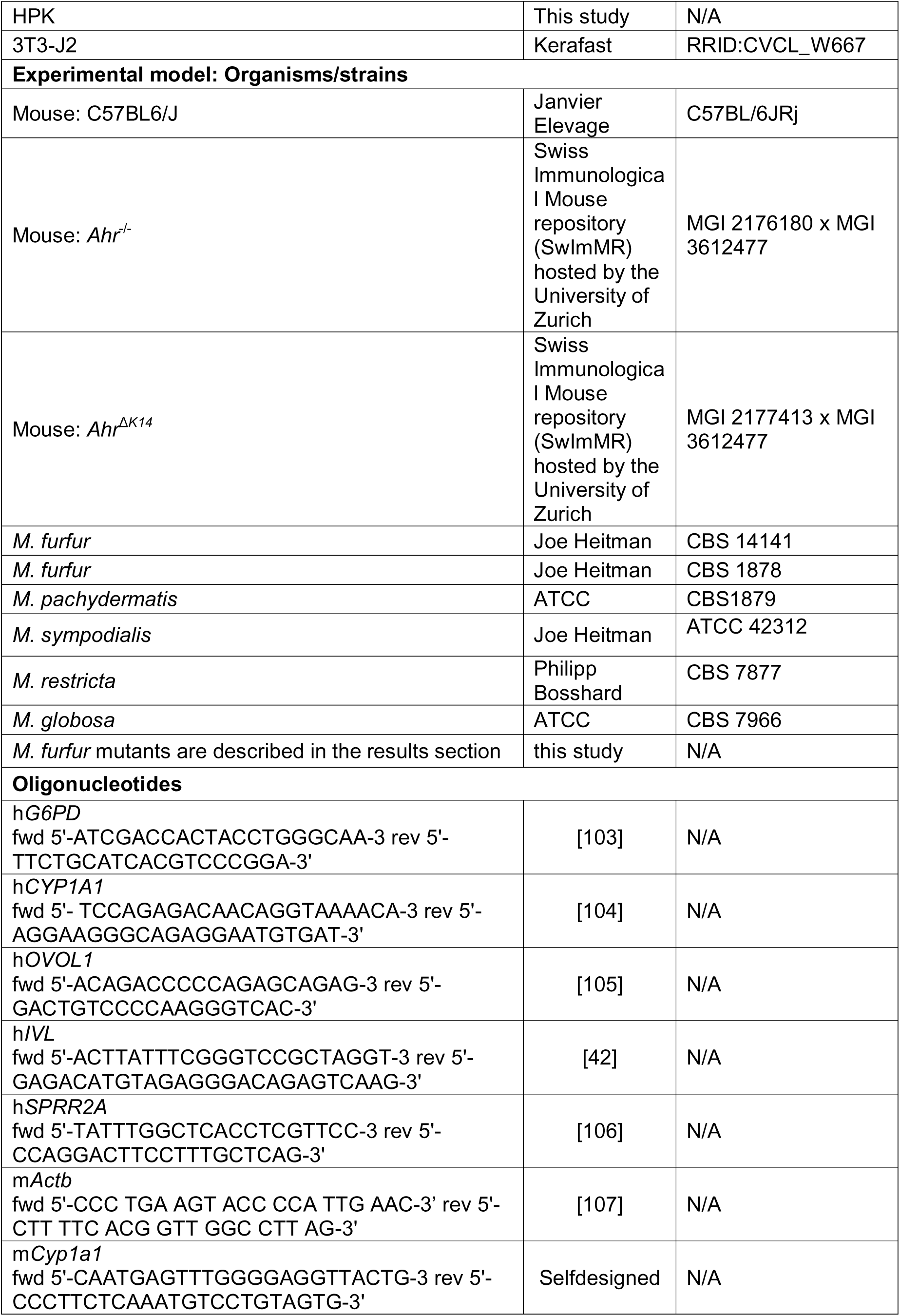
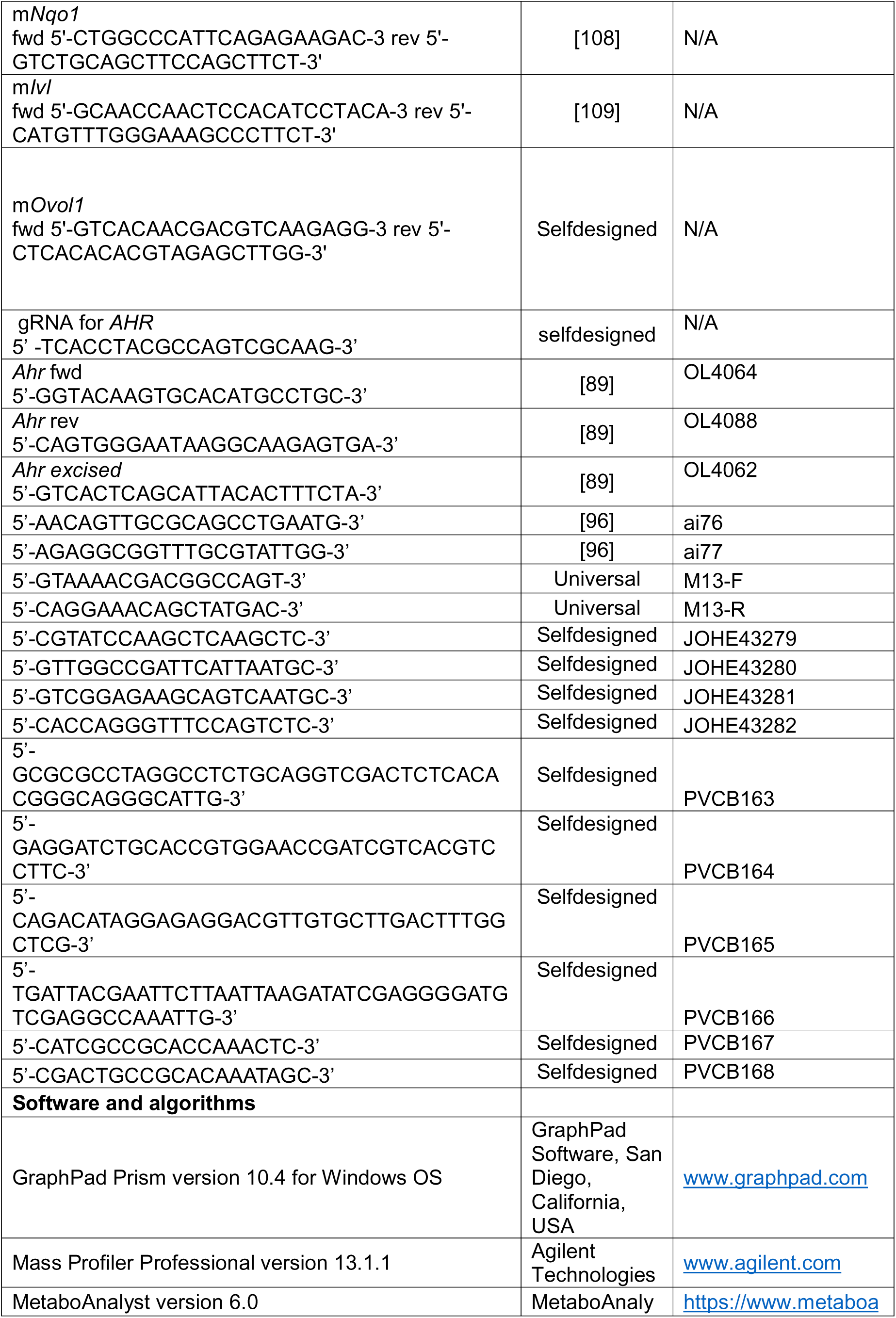
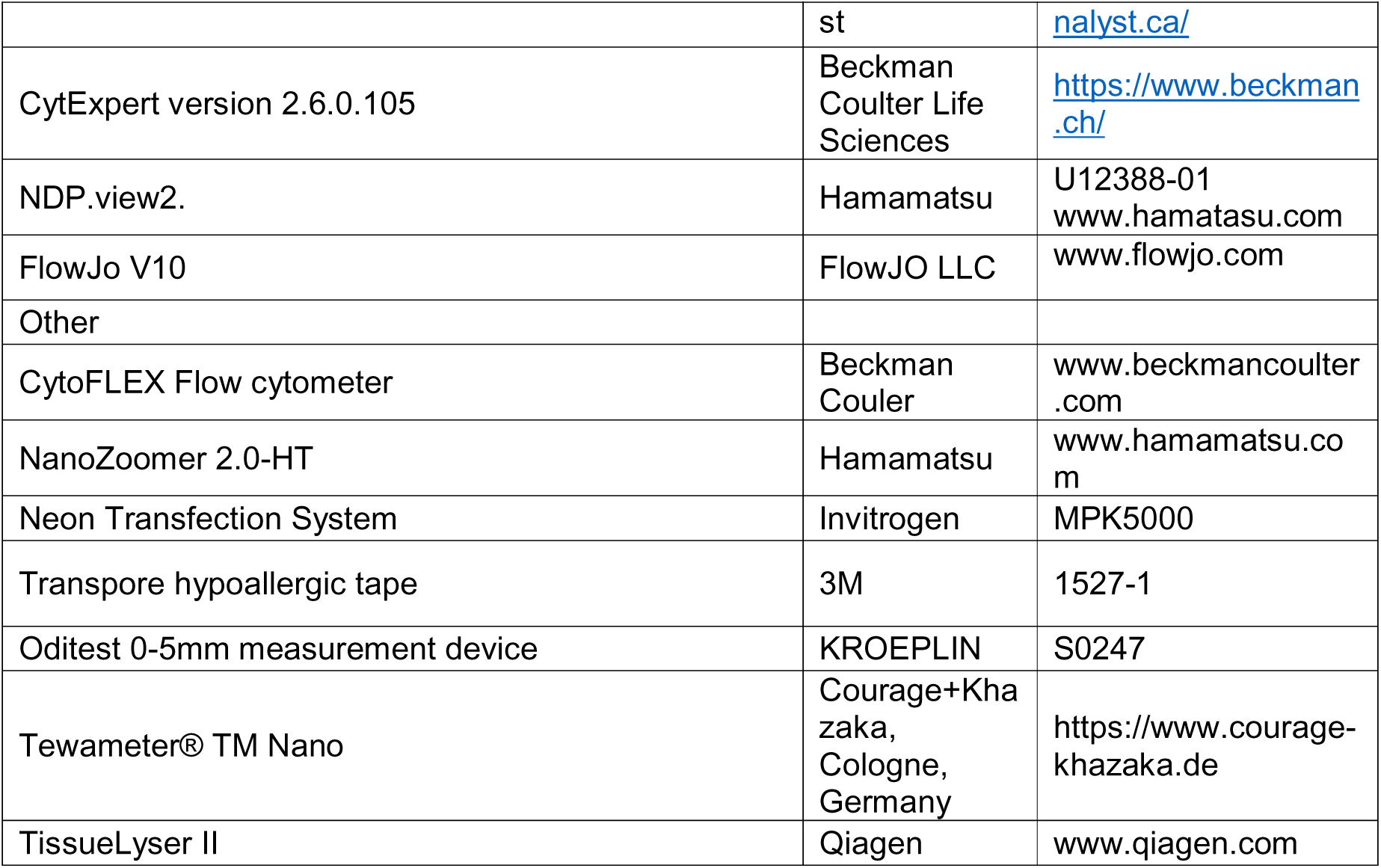

### Contact for Reagent and Resource Sharing

Further information and requests for resources and reagents should be directed to and will be fulfilled, if reasonable, by the lead contact Salomé LeibundGut-Landmann.

## DATA AVAILABILITY STATEMENT

All raw data linked to this study will be made publicly available at zenodo.org upon acceptance of the manuscript (doi will be provided).

## ACKNOWLEDGMENTS

The authors would like to thank M. Denison for providing the H1G1-1C3 reporter cell line; the staff of the Laboratory Animal Service Center of University of Zürich for animal husbandry; staff of the Laboratory for Animal Model Pathology of University of Zürich for histology; Center for Clinical Studies (ZKS) for access to equipment; members of the LeibundGut and Ianiri labs for helpful advice and discussions. This work was supported by the LEO Foundation (grant # LF-OC-22_001060 to S.L.L. and G.I.) and the Swiss National Science Foundation (grant # 310030_189255 to S.L.L.).

## AUTHOR CONTRIBUTIONS

E.G.-I., G.I. and S.L.L. designed the study and wrote the manuscript. E.G.-I. performed most experiments and analyzed the data. M.S., G.B, T.K. and D.B. performed selected experiments. A.S. and F.V. conducted metabolomics. S.L.L. and G.I. oversaw the study design and data analysis and acquired funding. All authors discussed the results and commented on the manuscript.

## SUPPLEMENTARY FIGURE LEGENDS

Supplementary Figure S1 (related to Figure 1). M. furfur-derived metabolites induces strong AhR activation via production of indoles in a tryptophan dependent manner.

**A.** *M. furfur* strains CBS1871 and CB14141 grown for 8 days on Tween 80 agar supplemented with 0, 0.1, 0.25 and 0.5 mg/mL L-tryptophan.

**B. – C.** *CYP1A1* expression by human primary keratinocytes (HPK) after 24 hours of infection with *M. furfur* cells (B) or conditioned medium from *M. furfur* (C), grown in presence (+) or absence (-) of tryptophan. n = 6 / group pooled from 2 independent experiments. The mean of each group is indicated.

**C.** GFP expression by the reporter cell line H1G1-1C3 in response to 24 hours of stimulation with conditioned medium from the indicated *Malassezia* spp. grown without tryptophan (white bars) or in presence of 0.5 (grey bars) or 5 mg/ml tryptophan (black bars), respectively. Expression levels are displayed relative to unstimulated and FICZ-stimulated cells. Bars are the mean + SEM of 16 separately stimulated wells from 2 independent experiments.

**D.** Representative images of *Malassezia* spp. grown on Tween 80 agar plates supplemented with 0, 0.5 and 5 mg/ml L-tryptophan.

The statistical significance of differences between groups was determined by unpaired *t*-test (B, C) or by two-way ANOVA (D), * p < 0.05, ** p < 0.01, *** p < 0.001, **** p < 0.0001.

**Supplementary Figure S2 (related to Figure 2). Activation of *AhR* by *M. furfur* in the epidermis modulates the expression of structural protein encoding genes.**

**A. - D.** *CYP1A1* (A, B) and *OVOL1*, *IVL*, and *SPRR2A* expression (C, D) by WT HEEs after 24 hours of infection with *M. furfur* (A, C) or exposure to conditioned medium from *M. furfur* (B, D) grown with (filled black symbols) or without tryptophan (filled grey symbols) or left uninfected (open symbols). n = 4 - 9 / group. Data in A - C are pooled from 2 independent experiments, data in D are from one experiment.

**A. E.** *CYP1A1* expression by two independent clones of *AHR* KO N/TERT1 and by parental WT N/TERT1 keratinocytes after 24 hours of stimulation with FICZ (filled symbols) or treatment with the solvent control (open symbols). n = 2 – 3 / group.

The mean of each group is indicated. The statistical significance of differences between groups was determined by one-way ANOVA (A - D) or two-way ANOVA (E), * p < 0.05, ** p < 0.01, *** p < 0.001, **** p < 0.0001.

**Supplementary Figure S3 (related to Figure 3). *M. furfur* induced AhR signaling restores barrier integrity in inflamed skin.**

**A.** The ear skin of C57BL/6 WT mice was mildly tape stripped prior to epicutaneously administering *M. furfur*, which was pre-cultured in presence (+Trp) or absence of tryptophan (-Trp). *Cyp1a1* relative expression was quantified from the epidermis of the colonized skin.

**B.** The ear skin of *Ahr*^-/-^ and *AhR^+/-^* littermates control mice was mildly tape stripped prior to epicutaneously administering *M. furfur*, which was pre-cultured in presence (+Trp) or absence of tryptophan (-Trp). Fungal load (CFU) was assessed on day 7 after fungal association. n = 7 – 9 / group, pooled from 2 independent experiments. The mean of each group is indicated.

**C.** Gating strategy for ear skin neutrophils quantified by flow cytometry in Figure 3D-E.

The statistical significance of differences between groups was determined by unpaired t test (A) or two-way ANOVA (B), * p < 0.05, ** p < 0.01, *** p < 0.001, **** p < 0.0001.

**Supplementary Figure S4 (related to Figure 4). AhR-mediated skin barrier restoration in response to *M. furfur* is keratinocyte intrinsic.**

**A.** AhR deletion in CD45^+^ and CD45-EpCam^+^ populations sorted from epidermal sheets of naïve *AhR*^Δ*K14*^ and *AhR^fl/fl^* littermate control mice.

**B.** The ear skin of *AhR*^Δ*K14*^ mice and *AhR^fl/fl^* littermate control mice was mildly tape stripped prior to epicutaneously administering *M. furfur*, which was pre-cultured in presence (+Trp) or absence of tryptophan (-Trp). A group of control mice was tape stripped and treated with olive oil (vehicle). Fungal load (CFU) was assessed on day 7 after fungal association. n = 8 / group pooled from 2 independent experiments, except for the vehicle group where n = 4-5. The mean of each group is indicated.

**Supplementary Figure S5 (related to Figure 5).**

**A.** *M. furfur* CBS 14141 grown in liquid mDix-mp supplemented with tryptophan (Trp), cystein (Cys) or methionine (Met) for 4 days.

**Supplementary Table S1**

Differentially produced by *M. furfur* CBS 14141 supplemented with tryptophan relative to metabolites produced without tryptophan supplementation.

**Supplementary Table S2.**

Differentially produced metabolites by *M. furfur sul1*Δ supplemented with tryptophan relative to *M. furfur* WT supplemented with tryptophan.

**Supplementary Table S3.**

Differentially produced metabolites by *M. furfur sul1*Δ supplemented with tryptophan relative to *sul1*Δ without tryptophan metabolism.

**Supplementary Table S4.**

Differentially produced metabolites by *M. furfur sul1*Δ relative to *M. furfur* WT.

