## Supplementary figures and images for "The skin commensal yeast *Malassezia* promotes tissue homeostasis via the aryl hydrocarbon receptor"

### Supplementary Figures S1 - S5

Suppl. Figure S1

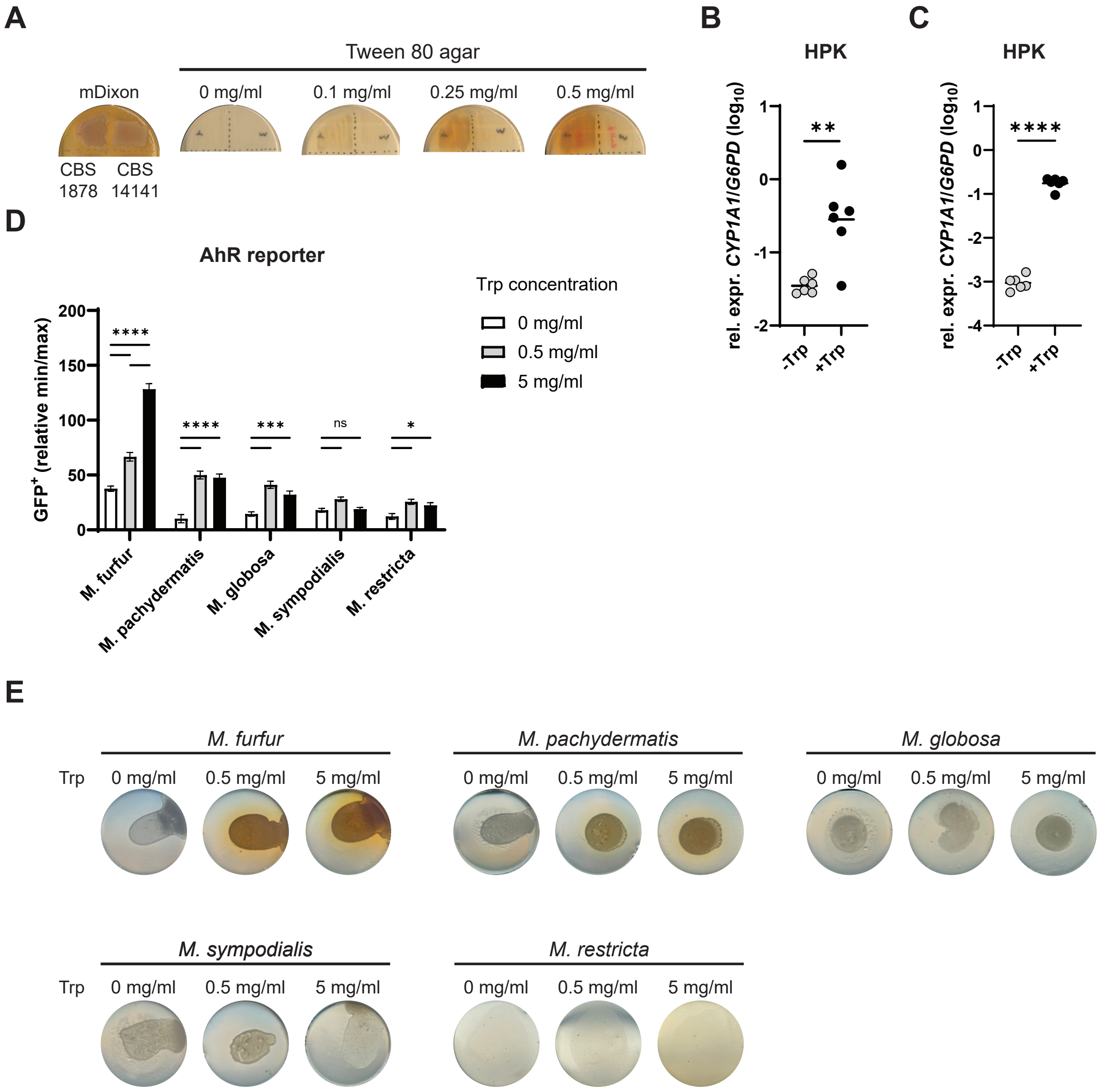

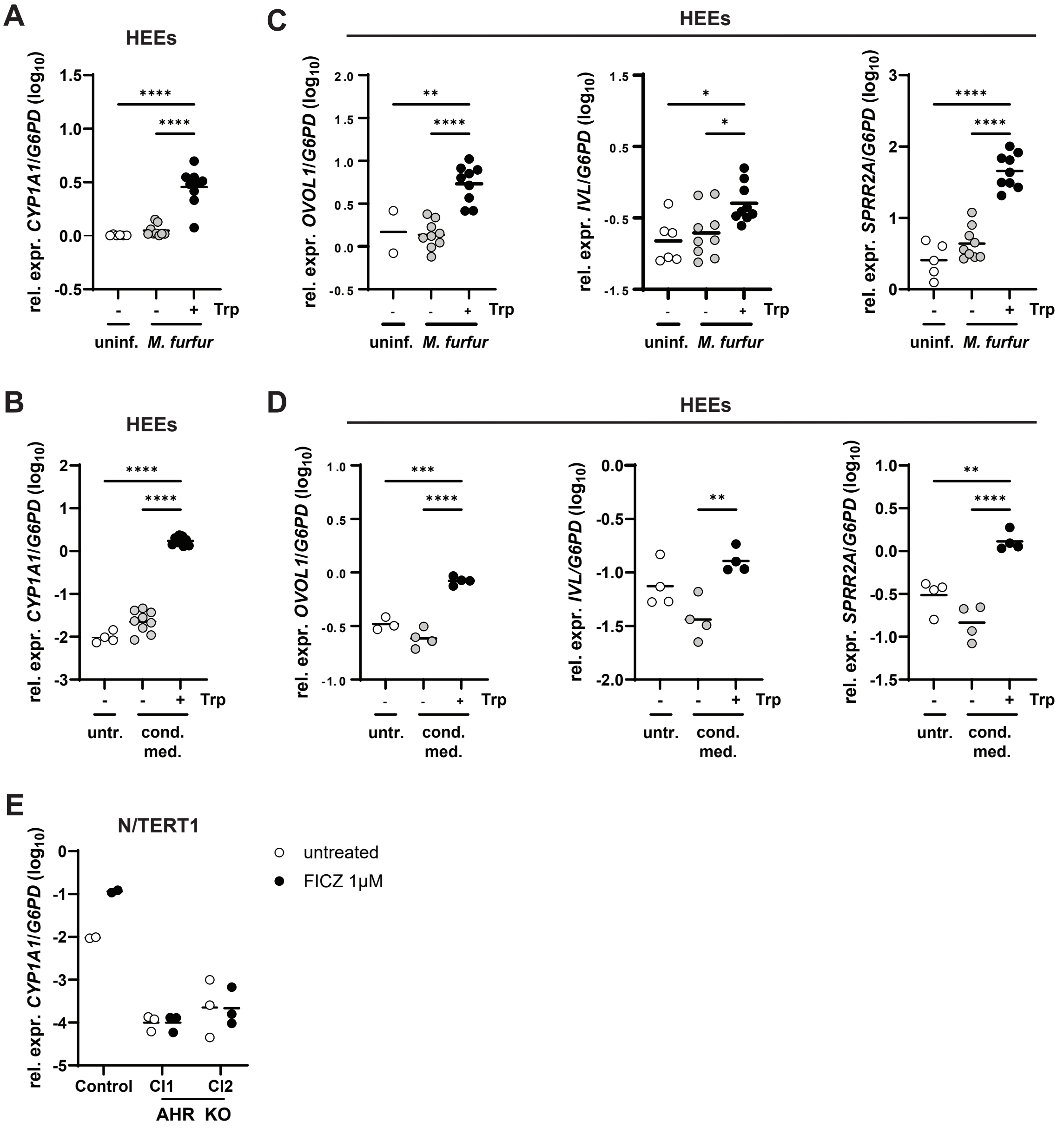

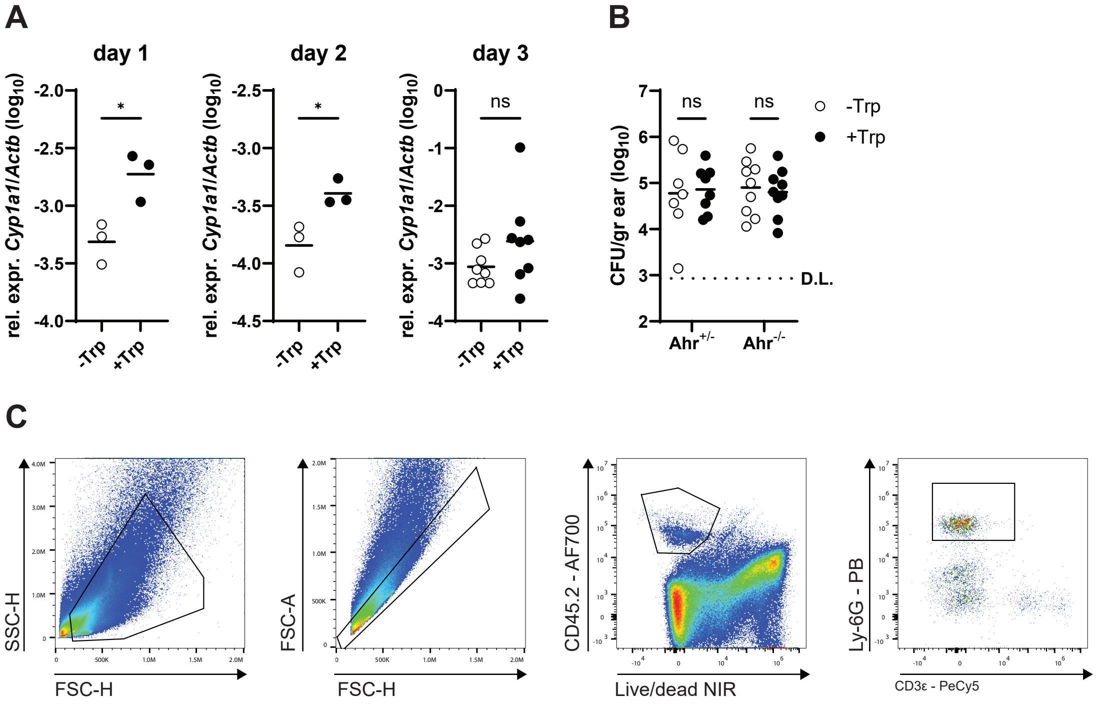

Suppl. Figure S4

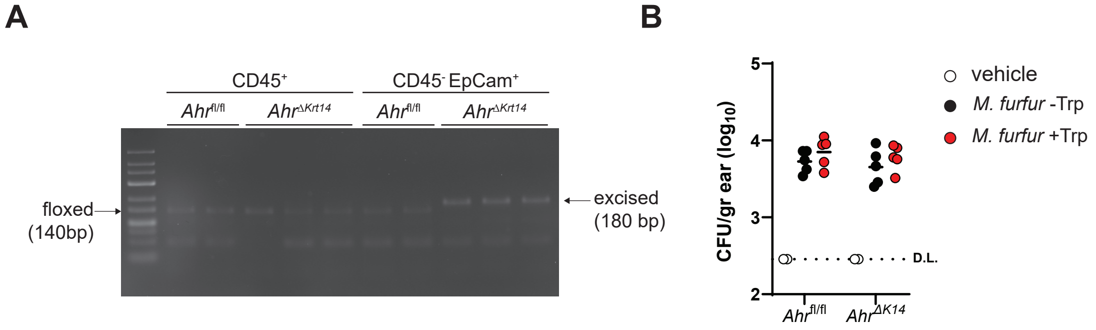

A

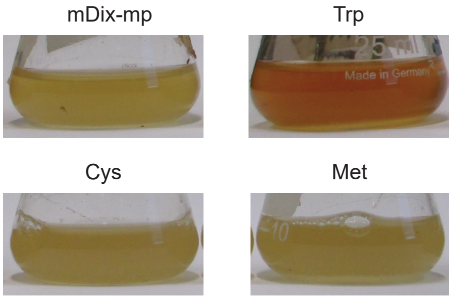
